# Molecular profiling of innate immune response mechanisms in ventilator-associated pneumonia

**DOI:** 10.1101/2020.01.08.899294

**Authors:** Khyatiben V. Pathak, Marissa I. McGilvrey, Charles K. Hu, Krystine Garcia- Mansfield, Karen Lewandoski, Zahra Eftekhari, Yate-Ching Yuan, Frederic Zenhausern, Emmanuel Menashi, Patrick Pirrotte

**Affiliations:** Collaborative Center for Translational Mass Spectrometry, Translational Genomics Research Institute, Phoenix, Arizona 85004; Translational Genomics Research Institute, Phoenix, Arizona 85004; HonorHealth Clinical Research Institute, Scottsdale, Arizona 85258; Applied AI and Data Science, City of Hope Medical Center, Duarte, California 91010; Center for Informatics, City of Hope Medical Center, Duarte, California 91010; Center for Applied NanoBioscience and Medicine, University of Arizona, Phoenix, Arizona 85004

**Keywords:** ventilator-associated pneumonia, endotracheal aspirate, proteome, metabolome, neutrophil degranulation

## Abstract

Ventilator-associated pneumonia (VAP) is a common hospital-acquired infection, leading to high morbidity and mortality. Currently, bronchoalveolar lavage (BAL) is utilized in hospitals for VAP diagnosis and guiding treatment options. While BAL collection procedures are invasive, alternatives such as endotracheal aspirates (ETA) may be of diagnostic value, however, their utility has not been thoroughly explored. Longitudinal ETA and BAL were collected from 16 intubated patients up to 15 days, of which 11 developed VAP. We conducted a comprehensive LC-MS/MS based proteome and metabolome characterization of longitudinal ETA and BAL to detect host and pathogen responses to VAP infection. We discovered a diverse ETA proteome of the upper airways reflective of a rich and dynamic host-microbe interface. Prior to VAP diagnosis by microbial cultures from BAL, patient ETA presented characteristic signatures of reactive oxygen species and neutrophil degranulation, indicative of neutrophil mediated pathogen processing as a key host response to the VAP infection. Along with an increase in amino acids, this is suggestive of extracellular membrane degradation resulting from proteolytic activity of neutrophil proteases. Days prior to VAP diagnosis, detection of pathogen peptides with species level specificity in ETA may increase specificity over culture-based diagnosis. Our findings suggest that ETA may provide early mechanistic insights into host-pathogen interactions associated with VAP infection and therefore facilitate its diagnosis and treatment.

**Graphical abstract:** 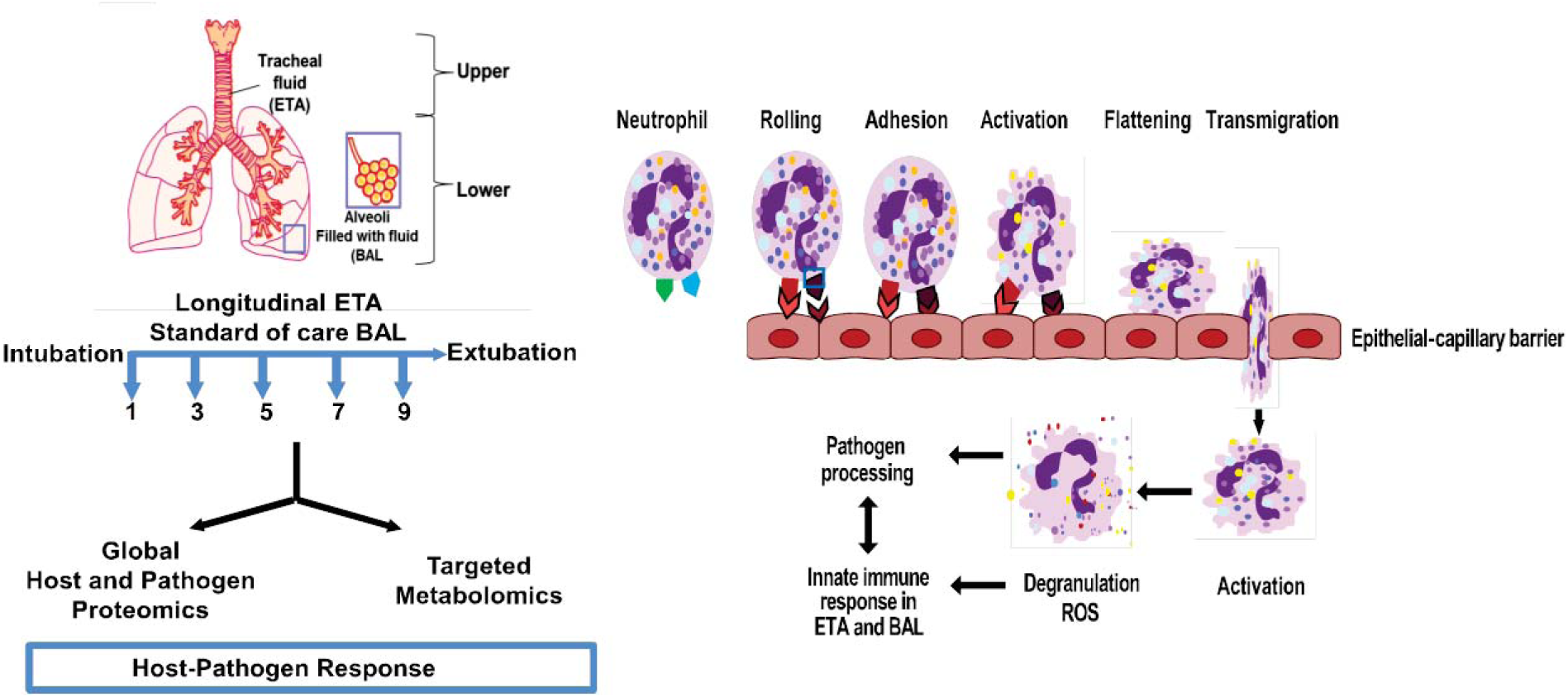

## Introduction

Ventilator-associated pneumonia (VAP) is the second most common hospital-acquired infection (HAI) in intensive care units (ICU), and is associated to 60% of all HAI-related deaths in the United States (1). It occurs 48 hours after mechanical ventilation and accounts for more than 300,000 cases annually in the US. (1, 2). Mechanical ventilation can injure tracheal epithelia, and promote environmental microbial colonization and migration from the upper to lower airways (3). All ICU patients with prolonged hospitalization are at high risk of developing VAP, increasing the cost of hospital stay by $40,000 to $50,000 per patient (3–5). VAP diagnosis is largely based on clinical criteria such as fever, infiltrate on chest radiograph, leukocyte counts, and positive cultures from bronchoalveolar lavages (BAL) (2, 6). With trauma patients, these symptoms are often non-specific and lead to overestimation of true VAP episodes resulting in the prescription of inadequate broad-spectrum antibiotics (6, 7). A study involving 300 U.S. hospitals showed a 52% increase in antibiotic consumption rate in intensive care unit (ICU) compared to non-critical care (8). In addition, VAP accounts for over half of the antibiotics used in the ICU which may lead to multi-drug resistance (9). Culture-based testing can reduce antibiotic misuse, but it is time-consuming and can delay diagnosis and treatment. Therefore, careful investigation of VAP pathogenesis is required to better understand host response to microbial dysbiosis and provide insights into the molecular mechanisms underlying the progression of infection.

Previous studies have proposed several candidate biomarkers (interleukin-1β, interleukin-8, soluble triggering receptor expressed on myeloid cells type 1, C-reactive protein (CRP), procalcitonin, and the mid-region fragment of pro-adrenomedullin) to assist VAP diagnosis in serum, plasma or BAL (10–13). Of these, interleukin-1β and interleukin-8 have been successfully validated in VAP diagnosis (13). Also, most of these proteins are inflammation markers and have shown variable sensitivity and specificity towards VAP detection (12, 14). This raises potential issues of misdiagnosis and emphasizes the need for further research on specific VAP biomarkers.

BAL has been a widely accepted matrix to study pulmonary infections (15, 16). Many studies have demonstrated the utility of BAL for microbial culture, 16S rDNA analysis and determining host-response against VAP infection (16–19). Endotracheal aspirate (ETA) is regarded as a source for non-invasive respiratory sampling and recently has been recommended for semi quantitative cultures in VAP diagnosis (20). However, the molecular composition of ETA has not been explored as extensively as BAL to understand host responses to infection. We hypothesize that reduced invasiveness involved in ETA sampling is permissive to more frequent longitudinal molecular snapshots of host immune response and changes to microbial flora during early infection. We anticipate that this enhanced granularity provides valuable mechanistic insights into VAP pathogenesis. We used a multi-disciplinary approach integrating proteomics and quantitative metabolomics on longitudinal ETA and matched BAL collected from intubated patients for this study.

## Experimental Procedures

### Chemicals

Chemicals and solvents were procured from Sigma-Aldrich (St. Lois, MO) or Fisher Scientific (San Jose, CA) unless otherwise stated. The chemicals used in this study were AR grade, and the formic acid (FA) and solvents were LC-MS grade.

### Specimen collections

Patients under mechanical ventilation at the ICU trauma center at HonorHealth Osborne Medical Center, Scottsdale, AZ were enrolled for this study. A written informed consent was obtained from either patient or a legal relative. The clinical protocol for sample collection was approved by the hospital’s Institutional Review Board and the Western Institutional Review Board, Puyallup, WA. All experimental procedures conformed to the principles set out in the Declaration of Helsinki and the Department of Health and Human Services Belmont Report. Patients with positive clinical symptoms (≥ 48 hours of intubation, fever (> 38.4 °C), increases in purulent secretions, new or progressive pulmonary infiltrates on chest radiograph), and a positive culture test using BAL for pathogenic microflora (*Escherichia coli*, *Pseudomonas aeruginosa*, *Staphylococcus aureus*, *Serratia marcescens*, *Streptococci* group C, *Enterobacter cloacae*, *Enterobacter aerogenes*, *Proteus mirabilis*, or *Candida albicans*) were diagnosed with VAP. The quantitative cultures were performed using BAL and two cut-offs, such as 10,000-100,000 CFU (colony forming units) for predominant signal pathogen and >100,000 CFU for 1-2 potential pathogens were employed to call the test positive. The chest x-ray was performed on all the patients enrolled in the study. The patients with no signs of clinical symptoms were referred as control. ETA was collected every other day, starting at the first day of intubation, until extubation. Upon positive clinical symptoms, BAL was collected as part of standard-of-care procedures (Figure 1) and used for microbial cultures to aid in clinical diagnosis. BAL collection was done by introducing 50 cc of sterile normal saline to the bronchial lumen to remove mucus plugs, secretions and debris. The detail description of study cohort and longitudinal collections are summarized in Table 1. Both the ETA and BAL biospecimens were collected in sterile tubes and immediately frozen and stored at −80 °C at the collection site. The samples were thawed, filter sterilized with 0.45 μm Ultrafree-CL HV centrifugal filters (Millipore, Billerica, MA), and stored at −80 °C until further processing. In this study, *Baseline* was defined as the first day of intubation for both control and VAP patients, and *VAP positive* as the day of VAP diagnosis. ETA collected two days before or after VAP diagnosis was defined as *pre-VAP* and *post-VAP*, respectively. Other time points in control groups were defined as *Control*. This classification was employed for both proteomics and metabolomics data analysis.

**Figure 1:**
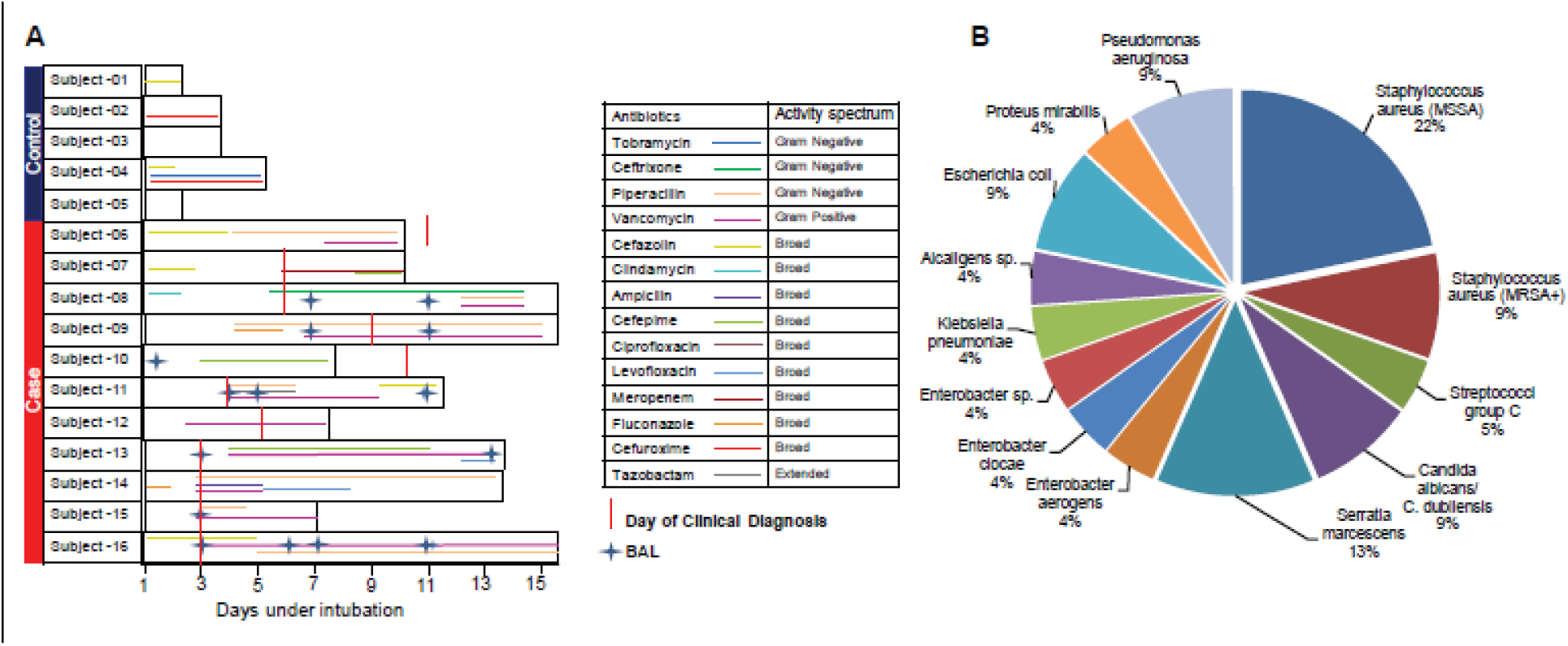
A. Patient cohort, sample collections and antibiotic treatment. Sixteen intubated patients enrolled in the study and were categorized in the control and case groups based on clinically diagnosed ventilator-associated pneumonia (VAP). The duration of intubation is denoted by the length of black outlined box for each patient, while the duration of antibiotic treatment is denoted by the length of colored line and the type of antibiotic treatment is provided in the legend. The endotracheal aspirate (ETA) samples were collected every other day throughout the intubation period. The bronchoalveolar lavage (BAL) samples were collected as indicated. **B.** Percentage of VAP pathogens detected in BAL culture in clinically diagnosed patients. MSSA = methicillin-sensitive *Staphylococcus aureus*, MRSA^+^ = methicillin-resistant *S. aureus*.

### Proteomics analysis

ETA and BAL were concentrated using Amicon ultra-3kDa centrifugal filters (Sigma-Aldrich, St Louis, MO) using the vendor’s protocol. The protein concentrations were measured using the Pierce BCA assay kit (ThermoFisher Scientific, San Jose, CA). Various methodologies for preparing BAL and ETA samples for shotgun proteomics analysis were evaluated and are summarized in Supplemental Methods S1. Equal protein amounts (160 μg) of each BAL or ETA sample were immunodepleted using Pierce Top 12 Abundant Protein Depletion spin columns (ThermoFisher, Scientific, San Jose, CA). The flow-through protein solution was subjected to a modified filter-aided sample preparation (21). Briefly, the samples were buffer exchanged to 50 mM ammonium bicarbonate buffer, pH 7.8, and the proteins were denatured using 8 M urea (1 hour), reduced using 1 mM dithiothreitol (1 hour, 37 °C), and alkylated with 40 mM iodoacetamide (1 hour, 37 °C). The proteins were digested with Trypsin Gold overnight at 37°C (Promega, Madison, WI). The resulting peptides were de-salted using C18 SPE cartridges (Waters, Milford, MA) and eluted with 70% acetonitrile (ACN) with 0.1% trifluoroacetic acid (*v/v*) (Waters). The eluted peptides were vacuum-dried and frozen at −20 °C until LC-MS/MS analysis. On the day of analysis, the samples were reconstituted in 0.1% formic acid. Sample preparation and data acquisition were randomized separately in order to minimize bias.

LC-MS/MS analysis was performed using a nanoAcquity ultra performance liquid chromatography system (Waters, Milford, MA) coupled to an Orbitrap Fusion Lumos Tribrid mass spectrometer (Thermo Fisher, San Jose, CA). One μg of peptides from each sample was separated on a BEH C18, 1.7 μM, 0.1 x 100 mm column (Waters, Milford, MA) using a 83.5 minute gradient from 3 to 90% solvent B (ACN, 0.1% FA) and 97 to 10% solvent A (Water, 0.1% FA) at a flow rate of 0.5 μL/min. The following gradient conditions were employed: 3 to 7% B for 1 minute, 7 to 25% B for 1 to 72 minutes, 25 to 45% B for 10 minutes, 45 to 90% B for 0.5 minutes, 90% B for 0.5 minute and column equilibration at 3% B for 10 minutes. MS spectra were acquired over a scan range of *m/*z 380 to 2000 using the orbitrap at 120,000 resolution followed by quadrupole isolation (width 1.6 TH) of precursor ions for data-dependent higher-energy collisional dissociation MS/MS with top speed, target automatic gain control values of 50,000, a 60 milliseconds maximum injection time and dynamic exclusion of 60 seconds. Precursors with charge states of +2 to +7 were fragmented at a normalized collision energy of 35% in the ion trap. *E. coli* tryptic digest (Waters, Milford, MA) (250 ng on column) were injected before and after each batch and brackets of 5 samples to assess inter-run variability. Raw data were searched with the Mascot search engine (MatrixScience, Boston, MA) against a recent human database (2015) on Proteome Discover Version 2.1.1.21 (Thermo Scientific) using the following parameters: trypsin rules, maximum 2 missed cleavages, fixed cysteine carbamidomethylation (+57.021 Da) and variable methionine oxidation (+15.995 Da). The precursor and product ion mass tolerances were set to 10 ppm and 0.5 Da, respectively. The target false discovery rate (FDR) was set to 1%. Peptide quantitation was performed using label-free quantitation based on precursor ion area under the curve (AUC).

For microbial proteomics, we curated a list of common VAP pathogens comprised of gram-positive, gram-negative bacteria and yeast (Supplemental Table S1) from literature. Proteome FASTA files of the listed pathogens were used as microbiome database and proteomics raw data generated in this study were searched with Mascot with above mentioned parameters. Uniqueness of peptide sequence of proteins towards pathogen genera and species were determined using blastP against a uniprotKB database. For validation of mass spectrometry observations, MPO (ab119605) and ELANE (ab11955) levels were measured by ELISA (Abcam, Cambridge, MA) in both VAP and control ETA samples according to instructions from the manufacturer.

**Table 1.**
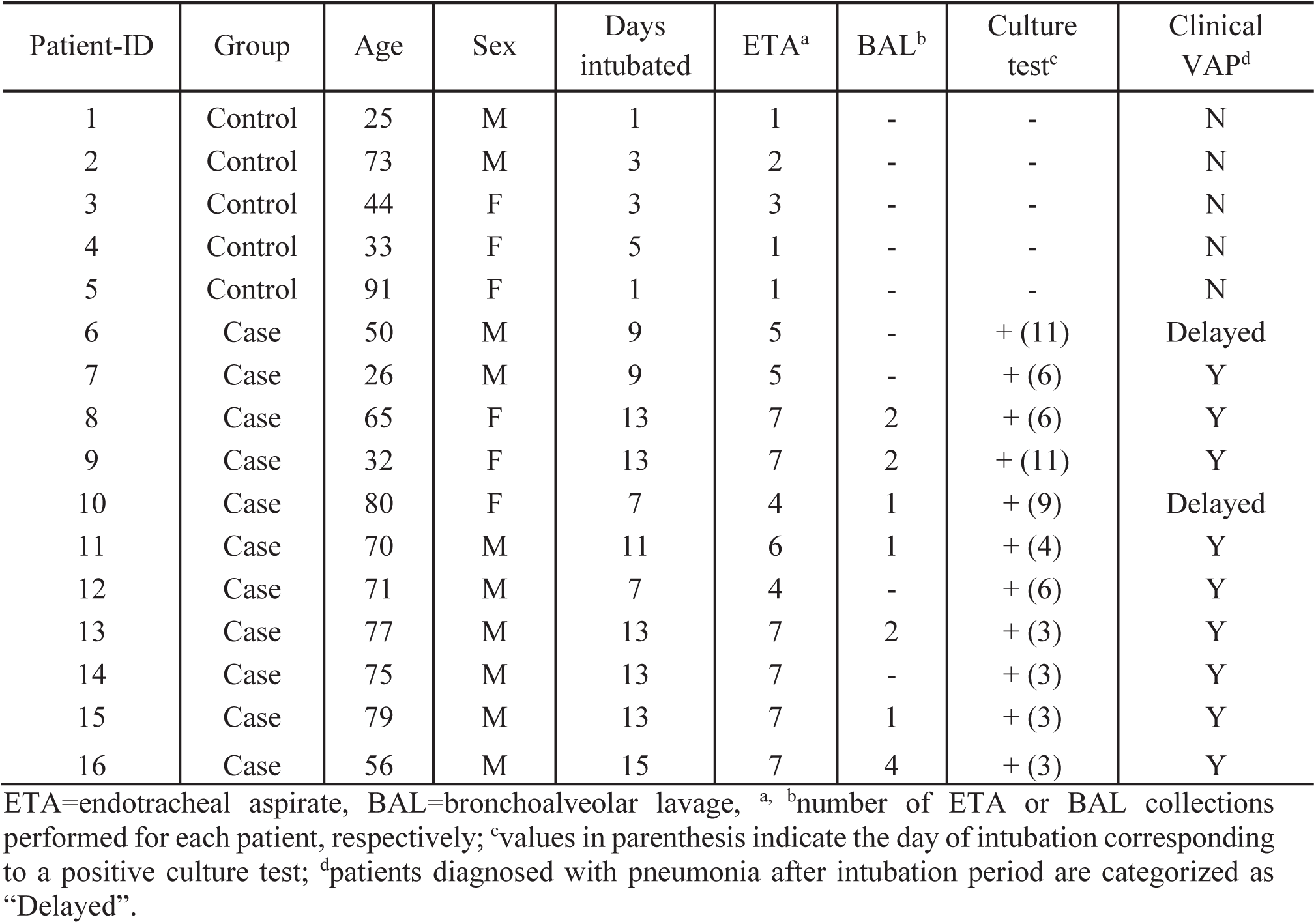
Cohort characteristics.

| Patient-ID | Group | Age | Sex | Days intubated | ETA<sup>a</sup> | BAL<sup>b</sup> | Culture test<sup>c</sup> | Clinical VAP<sup>d</sup> |
| --- | --- | --- | --- | --- | --- | --- | --- | --- |
| 1 | Control | 25 | M | 1 | 1 | - | - | N |
| 2 | Control | 73 | M | 3 | 2 | - | - | N |
| 3 | Control | 44 | F | 3 | 3 | - | - | N |
| 4 | Control | 33 | F | 5 | 1 | - | - | N |
| 5 | Control | 91 | F | 1 | 1 | - | - | N |
| 6 | Case | 50 | M | 9 | 5 | - | +(11) | Delayed |
| 7 | Case | 26 | M | 9 | 5 | - | +(6) | Y |
| 8 | Case | 65 | F | 13 | 7 | 2 | +(6) | Y |
| 9 | Case | 32 | F | 13 | 7 | 2 | +(11) | Y |
| 10 | Case | 80 | F | 7 | 4 | 1 | +(9) | Delayed |
| 11 | Case | 70 | M | 11 | 6 | 1 | +(4) | Y |
| 12 | Case | 71 | M | 7 | 4 | - | +(6) | Y |
| 13 | Case | 77 | M | 13 | 7 | 2 | +(3) | Y |
| 14 | Case | 75 | M | 13 | 7 | - | +(3) | Y |
| 15 | Case | 79 | M | 13 | 7 | 1 | +(3) | Y |
| 16 | Case | 56 | M | 15 | 7 | 4 | +(3) | Y |

ETA=endotracheal aspirate, BAL=bronchoalveolar lavage, <sup>a</sup>, <sup>b</sup>number of ETA or BAL collections performed for each patient, respectively; <sup>c</sup>values in parenthesis indicate the day of intubation corresponding to a positive culture test; <sup>d</sup>patients diagnosed with pneumonia after intubation period are categorized as “Delayed”.

### Targeted metabolomics analysis

The AbsoluteIDQ® p180 kit (Biocrates Life Sciences AG, Innsbruck Austria) was utilized to quantify 185 metabolites. Extractions and data analysis were performed using the vendor’s instructions. Briefly, 10 μL of BAL or ETA were subjected to phenylisothiocyanate derivatization followed by solvent extraction. A sample pool of ETA and BAL collections were prepared and used as sample quality control (QC). Triplicate QCs were distributed at equally through the samples. Data acquisition was conducted on an Acquity UPLC coupled with a Xevo TQ-S mass spectrometer (Waters, Milford, MA). For all ETA and BAL, simultaneous quantitation was performed for the following: 21 amino acids, 21 biogenic amines, 40 acylcarnitines, 89 glycerophospholipids. The data were analyzed using MetIDQ™. All the concentrations were measured in micromolar unit. Following nomenclatures, lysophosphatidyl glycerophospholipids (LysoPC x:y), glycerophospholipids (PC aa x:y and PC ae x:y) and Sphingolipids (SM x:y, SM[OH] x:y) were used throughout this manuscript. The x represents as a number of carbons in side chain and y denotes number of unsaturated fatty chains. For fatty acids, “aa” and “ae” represented fatty acids with glycerol moiety and fatty acid with fatty alcohol and glycerol, respectively.

### Experimental Design and Statistical Rationale

The experimental design comprised longitudinal ETA and BAL collections from 16 intubated patients (11 VAP patients, and 5 controls as detailed in Table 1). For the VAP group, 3 to 8 longitudinal ETA samples were collected per patient, whereas in the control group, up to 3 ETA samples were collected per patient (total of 74 ETA and 13 BAL samples). Due to sample limitations, technical replicates were not performed. As patients underwent standard of care, randomization was not performed during collection. All samples were randomized for processing and data acquisition. Relative protein abundances were log2 transformed. In VAP patients, only proteins present in 80% of either *Baseline* or *VAP positive* were selected for downstream proteomic analysis. Abundance of these proteins was also compared with *Baseline* from control patients. For quantitative metabolomics, only metabolites with <20% coefficient of variation (CV) in measurement of QC samples were included for down-stream analyses. Metabolites with >50% missing values were removed from the analysis and the remaining missing values were subjected to multivariate imputation by chained equations (MICE) (22). Temporal clustering was performed using median protein abundance or median concentration of metabolites. The distribution of proteomics and metabolomics data was tested using the Shapiro-Wilk test. Both datasets were not normally distributed (p>0.05) and therefore subjected to non-parametric analysis using the Wilcoxon rank-sum test. The p-values were adjusted using the Benjamini-Hochberg *post hoc* test. The following sample groups were compared: *VAP positive* to *Baseline*, *pre-VAP* to *Baseline* and *post-VAP* to *Baseline*. Proteins or metabolites with *p < 0.05 or adj. p < 0.05* were considered as significantly different. Gene Ontology (GO) annotation was performed using ToppFun (23). Pathway analysis was performed using Reactome and Ingenuity Pathway Analysis (IPA, QIAGEN Inc.) (24, 25). The *p-*values and fold-changes for ETA proteins were input into IPA and mapped against the human Ingenuity Knowledgebase with default values to uncover enriched pathways in VAP patients. The activation z-score was calculated by IPA software to determine positive or negative enrichment of pathways, diseases and biological functions as categories. The score predicts the increase or decrease in form of positive or negative z-score, respectively. The proteomics and metabolomics data were further assessed for similarity between ETA and BAL matrices using Bland-Altman analysis (26).

## Results

### Study cohort

Our study cohort was composed of 16 trauma patients intubated up to 15 days. Eleven of these patients exhibited symptoms of pneumonia (VAP patients) and five did not present any signs of pulmonary infection (control patients). The clinical annotation and antibiotic regimens are described in Table 1 and Figure 1A. Duration of intubation was longer in VAP (≥ 7 days) than control patients (≤ 5 days). A total of 8 patients including 3 control patients and 5 VAP patients were given broad spectrum antibiotics at intubation, whereas no antibiotics were given to the remaining 2 controls and 6 VAP patients. In VAP patients, antibiotic prophylaxis at the time of intubation did not show any better protection compared to no antibiotics; also, there was no clear-cut effect of antibiotic prophylaxis on the length of mechanical ventilation. Further antibiotic treatment was aligned as per BAL culture and clinical diagnosis. Based on BAL culture, *Staphylococcus aureus* was the most common VAP pathogen observed: 7 patients harbored methicillin-sensitive *S. aureus* (MSSA) and 2 patients harbored methicillin-resistant *S. aureus* (MRSA) (Figure 1B). As all patients in our study cohort were administered a standard-of-care regimen of antibiotics, their impact on the patient proteome and metabolome was not evaluated.

### ETA proteome reveals neutrophil mediated response in VAP

We identified a total of 3067 unique proteins in ETA collections across all patients and time points. We compared patient-matched ETA and BAL collected on the same day and identified 1811 and 1097 unique proteins in ETA and BAL collected on the same day. Of these, 975 proteins represented 88.9% of BAL proteome of this cohort and mapped to 187 significant reactome pathways. The top 10 mapped pathways were neutrophil degranulation, innate immune system, immune system, complement cascade, regulation of complement cascade, platelet activation, signaling and aggregation, platelet degranulation, regulation of insulin-like growth factor transport, post-translational protein phosphorylation and hemostasis (Supplemental Table S2). These suggest enrichment of proteins associated to host immunity in both ETA and BAL. Further, Bland-Altman comparison of these shared proteins showed that there was no significant bias between ETA and BAL as most of the data sits between the 95% confidence interval upper limit of 4.1 and the lower limit of −4.4. Figure 2B). The GO terms for the biological processes related to innate and humoral immunity were similarly enriched across both fluids (Figure 2C). This suggests similarities in proteome composition between BAL and ETA.

**Figure 2:**
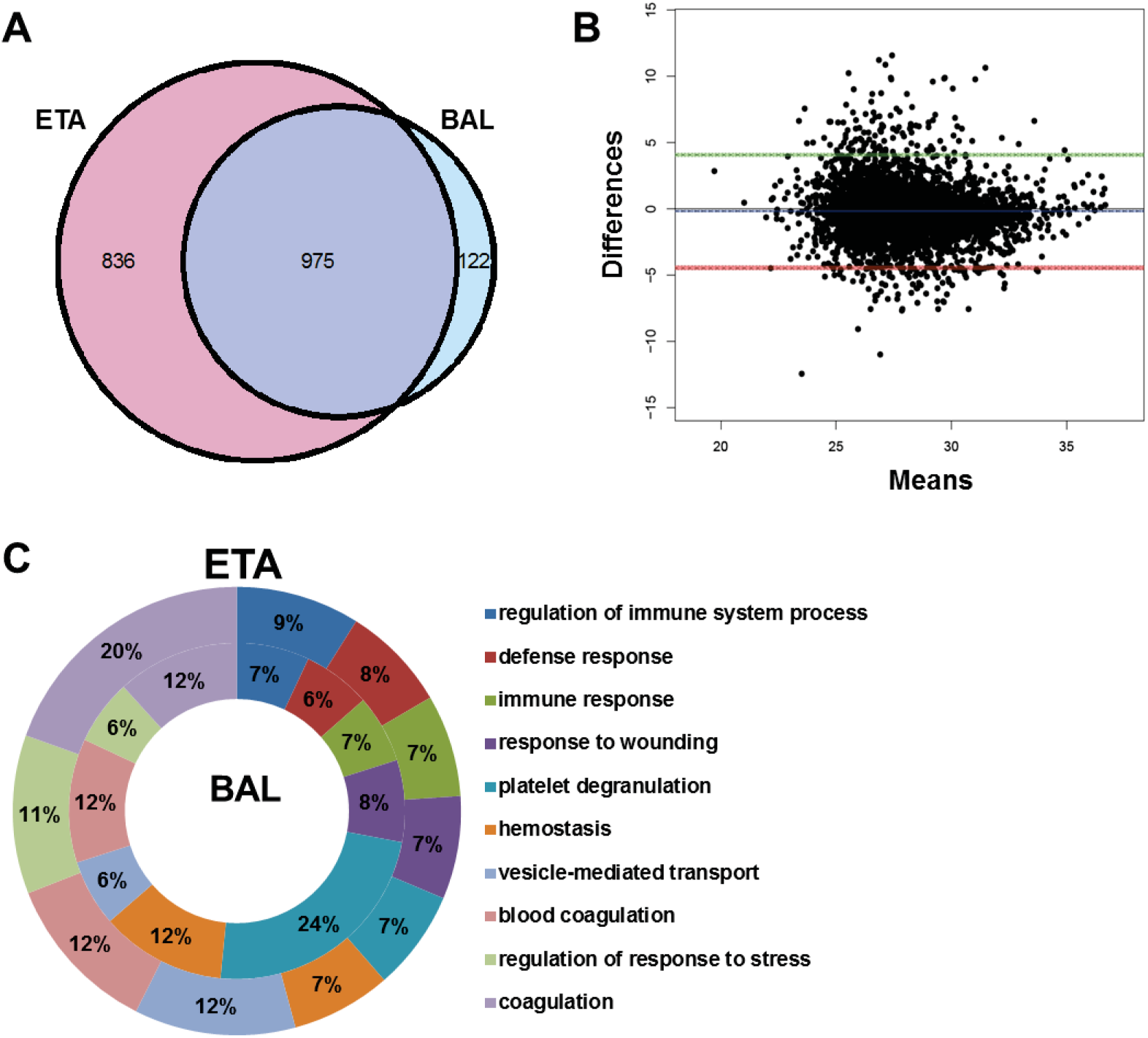
Comparison of ETA and BAL proteomes in VAP patients. A. The Venn diagram shows the numbers of proteins shared between ETA and BAL proteins. **B.** Bland-Altman analysis of bias and 95% limit of agreement for ETA versus BAL proteins**. C.** Gene ontology (GO) analysis (biological process) of ETA and BAL proteins. Doughnut plot shows the top 10 enriched GO terms of biological process for BAL (inner core) and ETA (outer core) proteins.

Due to the relative ease-of-access and increased sampling availability, we focused our study on ETA. Since intubation was variable across VAP patients and controls, we compared the first day of intubation (*Baseline*) against subsequent time points in VAP patients, and also compared *Baseline* ETA proteome in VAP and control patients.

The patients enrolled in our study were intubated for variable length of time and developed infection at different days. We categorized longitudinal collections to major clinical events of interest, such as *Baseline*, *VAP positive*, *pre-VAP* and *post-VAP*. Binning median protein abundance for each event, unsupervised temporal clustering showed concomitant clustering of *Baselines* of both control (Control *Baseline*) and VAP patients (VAP *Baseline*), suggesting that at the time of intubation, ETA proteomes of both control and VAP patients remained unchanged. (Figure 3A). The ETA proteome in VAP patients was following the pattern of progression of VAP infection from *pre-VAP, VAP positive* and *post-VAP* (Figure 3A).

**Figure 3.**
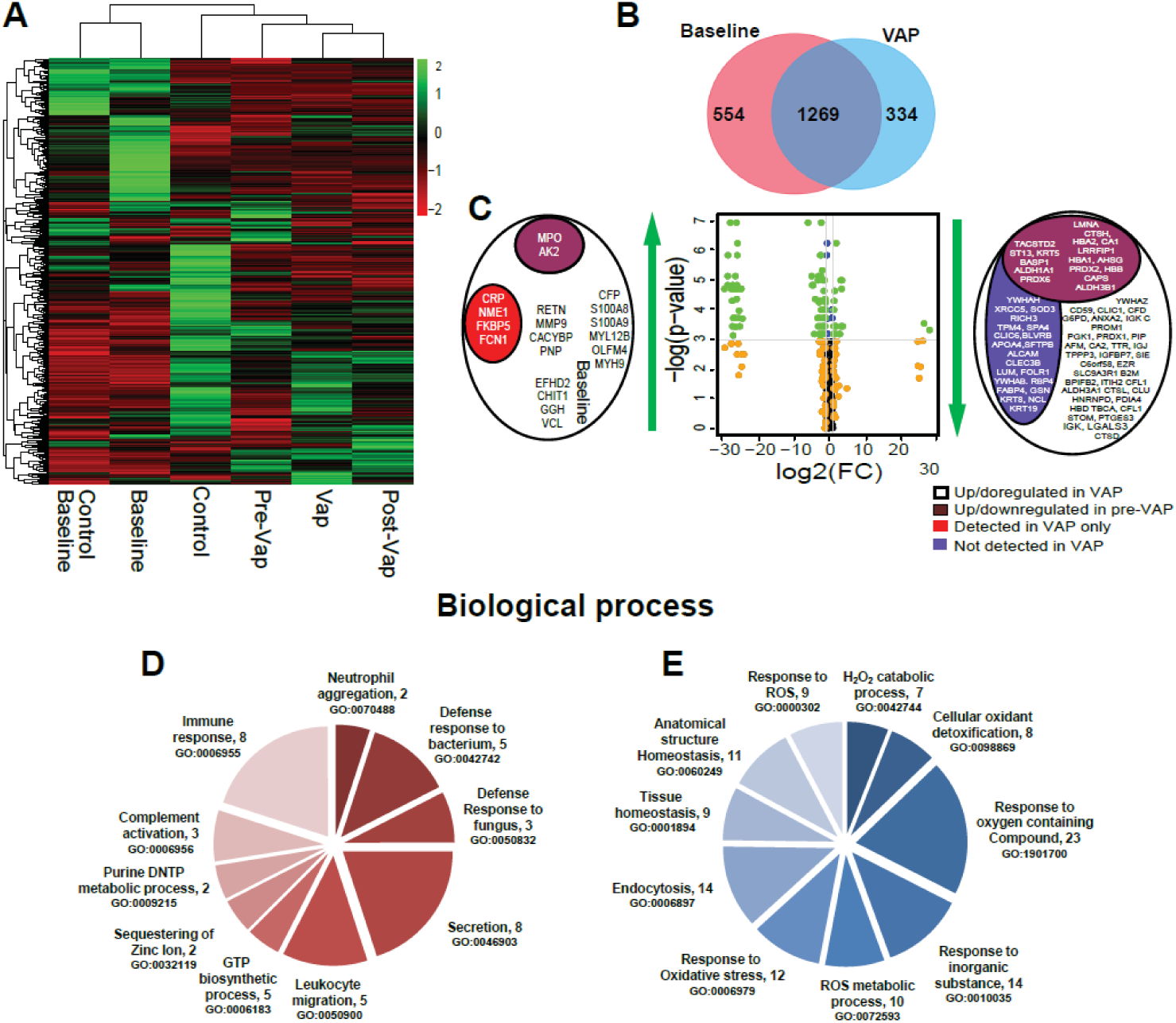
ETA proteome analysis. **A.** Temporal clustering of clinical time points using median protein abundance. **B.** Comparison of the *Baseline* and *VAP positive* ETA proteome in VAP patients. **C. Left panel**, significantly up-regulated proteins in VAP patients; **Center Panel**, Volcano plot representing the ETA proteome with, (i) in blue, significant proteins (*p* < 0.05, Wilcoxon rank-sum test), (ii) in orange, proteins for which the log2(FC) of the abundance (*VAP positive* compared to *Baseline*) > 1 or < −1, and (iii) in green, significant proteins (*p* < 0.05) with log2(FC) > 1 or < −1. **Right panel**, significantly down-regulated proteins. For the left and right panels, (i) in pink, a subset of up-regulated (left) and down-regulated (right) proteins in both *VAP positive* and *pre-VAP*, (ii) in red, proteins present in only *VAP positive* and absent in *Baseline*, and (iii) in purple, proteins absent in *VAP positive*. The list of proteins in left and right panels indicates significantly up- and down-regulated proteins, respectively. FC= Fold change. **D.** Gene Ontology (GO)-based functional enrichment analysis of significantly differentially abundant proteins in *VAP positive* compared to *Baseline* (biological process). Up-regulated (left) and down-regulated (right) GO categories (parenthesis). *Baseline* = day 1 of intubation for Control and VAP, *VAP positive =* day of VAP diagnosis*, pre-VAP =* a day before VAP, *post-VAP =* a day after VAP, *Control* = other intubation time points in control patients (day3 and day5)

In VAP patients, a total of 1823 and 1603 proteins were identified in the *Baseline* and *VAP positive* ETA, respectively, with 1269 proteins shared between time points (Figure 3B). Out of those, 10 to 19% of unique proteins were identified across all ETA, reflecting proteome variability in longitudinal samples.

The 1269 shared ETA proteins were used to compare the *Baseline* and *VAP positive* in VAP patients. Changes were further compared with *Pre-VAP* and *Post-VAP* groups. IPA analysis on all proteins between *VAP positive* and *Baseline* revealed 133 enriched pathways. *Complement system*, *Acute phase response signaling*, *Actin cytoskeleton signaling*, *Gluconeogenesis I*, *Integrin signaling*, *Glycolysis I*, *Regulation of actin-based motility by Rho*, and *Pentose phosphate pathway* are positively enriched, while *LXR/RXR activation* and *Remodeling of epithelial adherens junctions* are negatively enriched (Supplemental Table S3). GO terms above were further verified by the biological functions and diseases from IPA, which reported *degranulation of neutrophils, granulocytes,* and *phagocytes*, *leukocyte migration*, *inflammation*, *apoptosis*, and *necrosis* as most enriched (Supplemental Table S4). These pathways and processes were also observed to be enriched in *Pre-VAP* samples compared to *Baseline* and suggesting early activation of leukocyte mediated immunity in response to VAP pathogen.

Further statistical comparisons identified 96 differentially abundant proteins (*p* < 0.05, median fold change >2) between *Baseline* and *VAP positive* (Figure 3C) (Table 1). Twenty up-regulated proteins contributed to the following GO terms (biological processes, *p <* 0.05): *neutrophil aggregation*, *defense response to bacterium and fungus*, *leukocyte migration*, and *complement activation* (Figure 3D); 76 down-regulated proteins were linked to *reactive oxygen species metabolic process*, *oxidative stress*, *cellular oxidant detoxification*, and *tissue homeostasis* (Figure 3E). This suggests neutrophil mediated innate immune response and wound healing processes in *VAP positive*.

To determine response specificity against infection, we compared *Baseline* ETA between VAP and control patients. None of the 96 differentially abundant proteins identified from longitudinal analysis were significantly different across *Baselines.* This further corroborates our observation from temporal clustering. However, 26 proteins were not detected in *VAP positive* but were present in all *Baseline* (Figure 3B, Table 3). Functional annotation reveals their role in multiple binding activities (*p* < 0.05) (*hormone binding*, *vitamin binding*, *copper ion binding*, *scaffold protein binding*) (Figure 3A, Supplemental Table S5). Absence of these proteins in *VAP positive* may imply pathogen binding and clearance. Both isoform 2 of nucleoside diphosphate kinase A (NME-1) and CRP were detected in VAP ETA only (Table 3). To gain further insight, the significant differentially abundant proteins were mapped to Reactome pathways (Table 4). Two of the pathways with low *p-*values (*p* < 6.6E-14), *neutrophil degranulation* (11 proteins) and *innate immune system* (13 proteins) represent 55 to 65% of all up-regulated proteins, suggesting increased secretion of host immune proteins in VAP patients. Out of 76 down-regulated proteins (Figure 3C), 48 proteins were mapped to top 10 significant (*p* < 0.05) pathways, *erythrocytes take up oxygen and release carbon dioxide*, *neutrophil degranulation*, *Chk1/Chk2(Cds1) mediated inactivation of Cyclin B:Cdk1 complex*, *detoxification of reactive oxygen species*, *TP53 regulates metabolic genes*, *neurodegenerative diseases*, *scavenging of heme from plasma*, *amyloid fiber formation,* and *platelet degranulation.* Majority of up-regulated proteins mapped to multiple pathways linked to pathogen recognition and innate immunity, while most down-regulated blood proteins, carbonic anhydrase 1 (CA1), carbonic anhydrase 2 (CA2), hemoglobin subunit beta (HBB), hemoglobin subunit alpha (27), hemoglobin subunit delta (HBD), peroxiredoxin-1 (PRDX1), peroxiredoxin-2 (PRDX2), peroxiredoxin-6 (PRDX6) and erythrocyte band 7 integral membrane protein (STOM) mapped to tissue injury (Table 2, 4).

**Table 2.**
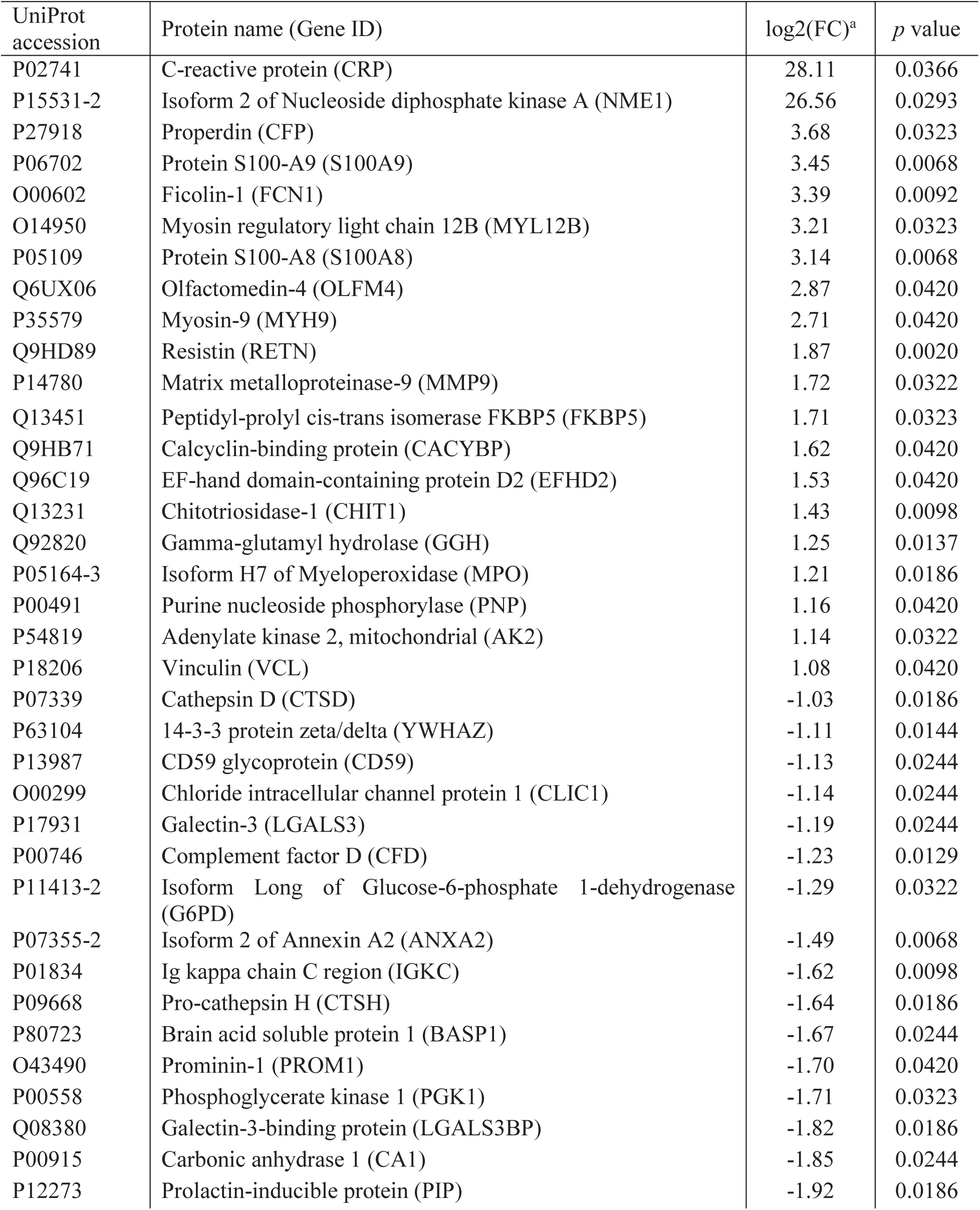
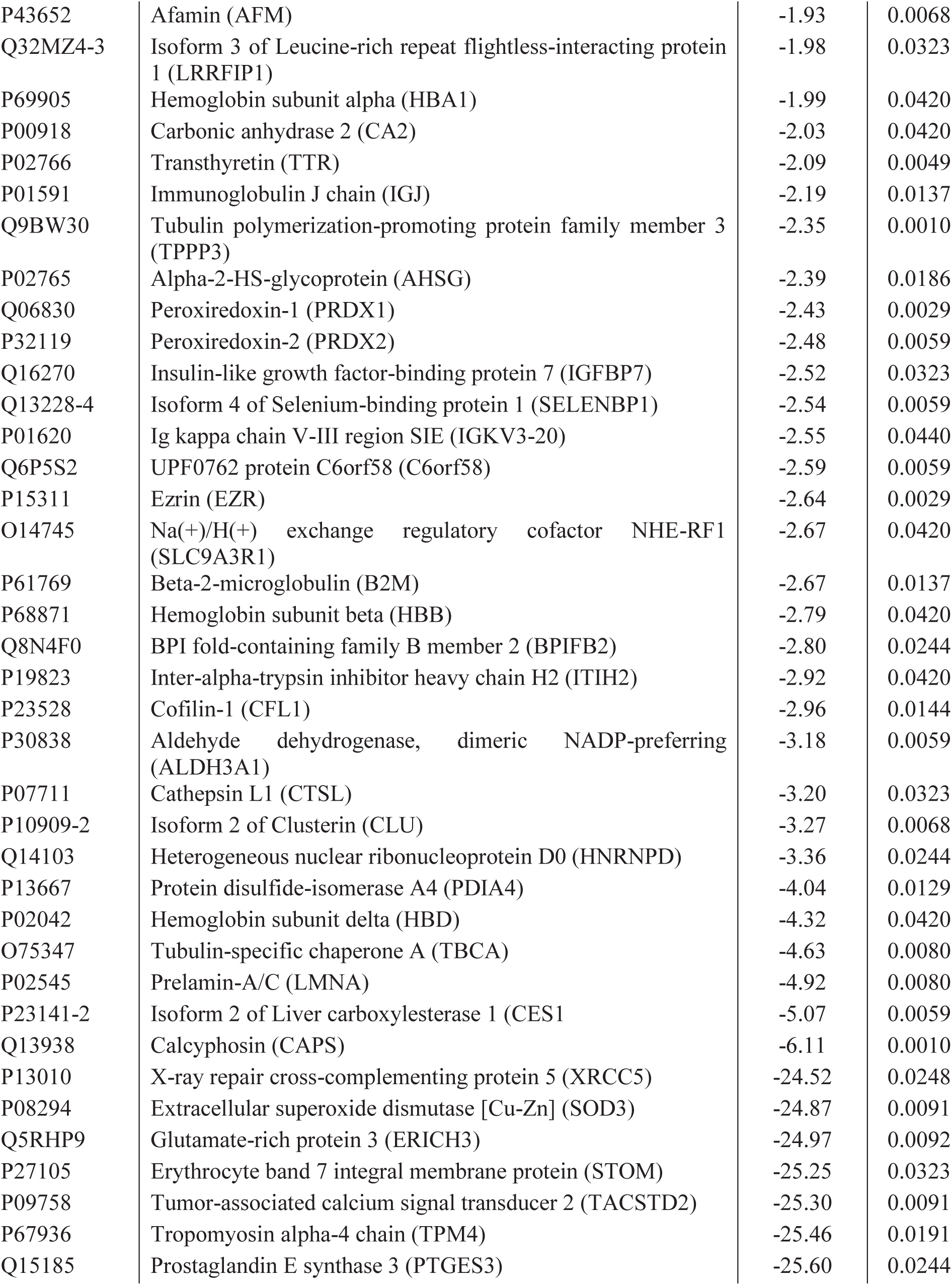
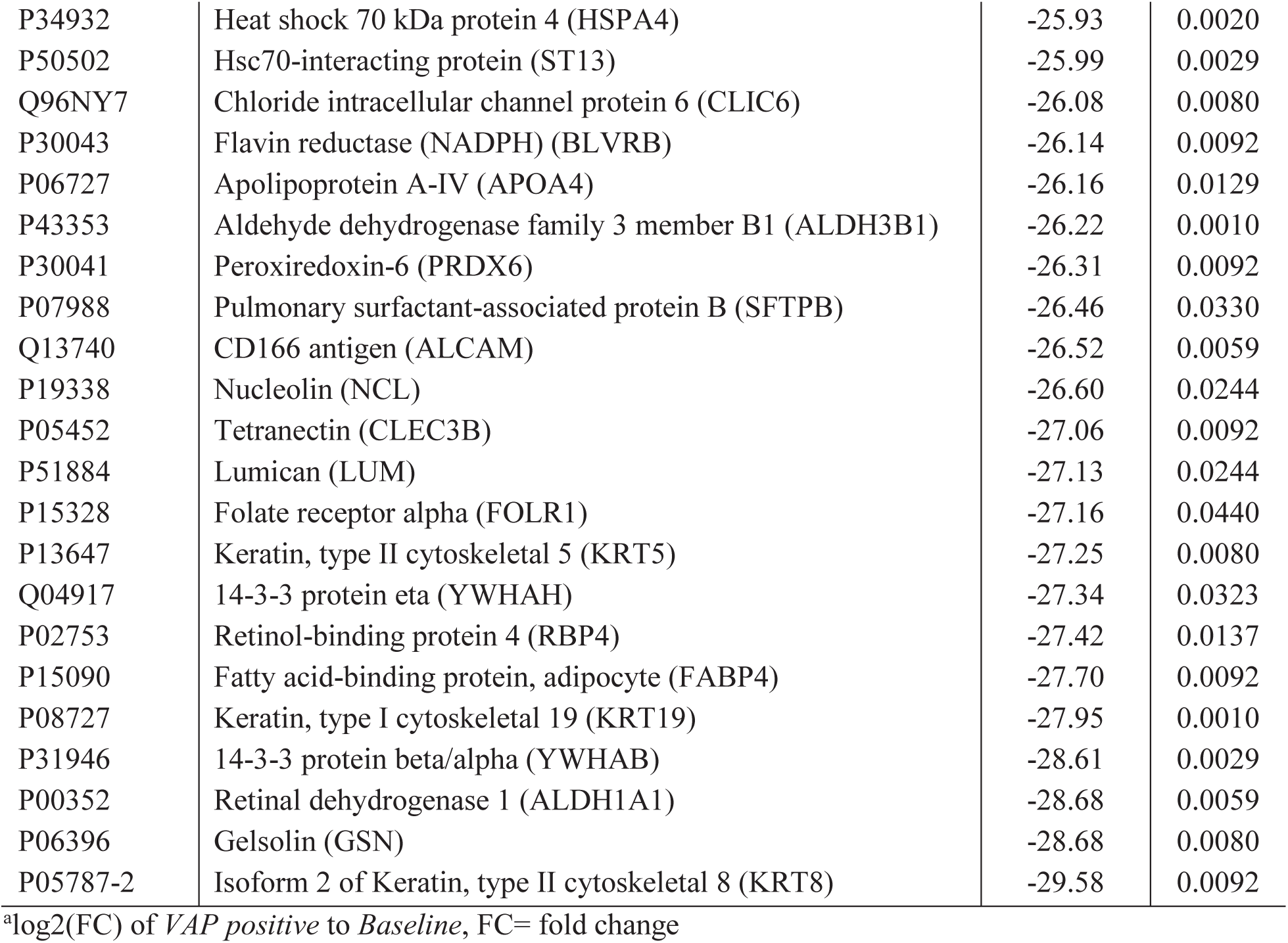
Differential expression of ETA proteins between Baseline and the day of clinical diagnosis (VAP positive)

| UniProt accession | Protein name (Gene ID) | log2(FC)<sup>a</sup> | p value |
| --- | --- | --- | --- |
| P02741 | C-reactive protein (CRP) | 28.11 | 0.0366 |
| P15531-2 | Isoform 2 of Nucleoside diphosphate kinase A (NME1) | 26.56 | 0.0293 |
| P27918 | Properdin (CFP) | 3.68 | 0.0323 |
| P06702 | Protein S100-A9 (S100A9) | 3.45 | 0.0068 |
| O00602 | Ficolin-1 (FCN1) | 3.39 | 0.0092 |
| O14950 | Myosin regulatory light chain 12B (MYL12B) | 3.21 | 0.0323 |
| P05109 | Protein S100-A8 (S100A8) | 3.14 | 0.0068 |
| Q6UX06 | Olfactomedin-4 (OLFM4) | 2.87 | 0.0420 |
| P35579 | Myosin-9 (MYH9) | 2.71 | 0.0420 |
| Q9HD89 | Resistin (RETN) | 1.87 | 0.0020 |
| P14780 | Matrix metalloproteinase-9 (MMP9) | 1.72 | 0.0322 |
| Q13451 | Peptidyl-prolyl cis-trans isomerase FKBP5 (FKBP5) | 1.71 | 0.0323 |
| Q9HB71 | Calcyclin-binding protein (CACYPB) | 1.62 | 0.0420 |
| Q96C19 | EF-hand domain-containing protein D2 (EFHD2) | 1.53 | 0.0420 |
| Q13231 | Chitotriosidase-1 (CHIT1) | 1.43 | 0.0098 |
| Q92820 | Gamma-glutamyl hydrolase (GGH) | 1.25 | 0.0137 |
| P05164-3 | Isoform H7 of Myeloperoxidase (MPO) | 1.21 | 0.0186 |
| P00491 | Purine nucleoside phosphorylase (PNP) | 1.16 | 0.0420 |
| P54819 | Adenylate kinase 2, mitochondrial (AK2) | 1.14 | 0.0322 |
| P18206 | Vinculin (VCL) | 1.08 | 0.0420 |
| P07339 | Cathepsin D (CTSD) | -1.03 | 0.0186 |
| P63104 | 14-3-3 protein zeta/delta (YWHAZ) | -1.11 | 0.0144 |
| P13987 | CD59 glycoprotein (CD59) | -1.13 | 0.0244 |
| O00299 | Chloride intracellular channel protein 1 (CLIC1) | -1.14 | 0.0244 |
| P17931 | Galectin-3 (LGALS3) | -1.19 | 0.0244 |
| P00746 | Complement factor D (CFD) | -1.23 | 0.0129 |
| P11413-2 | Isoform Long of Glucose-6-phosphate 1-dehydrogenase (G6PD) | -1.29 | 0.0322 |
| P07355-2 | Isoform 2 of Annexin A2 (ANXA2) | -1.49 | 0.0068 |
| P01834 | Ig kappa chain C region (IGKC) | -1.62 | 0.0098 |
| P09668 | Pro-cathepsin H (CTSH) | -1.64 | 0.0186 |
| P80723 | Brain acid soluble protein 1 (BASP1) | -1.67 | 0.0244 |
| O43490 | Prominin-1 (PROM1) | -1.70 | 0.0420 |
| P00558 | Phosphoglycerate kinase 1 (PGK1) | -1.71 | 0.0323 |
| Q08380 | Galectin-3-binding protein (LGALS3BP) | -1.82 | 0.0186 |
| P00915 | Carbonic anhydrase 1 (CA1) | -1.85 | 0.0244 |
| P12273 | Prolactin-inducible protein (PIP) | -1.92 | 0.0186 |
| P43652 | Afamin (AFM) | -1.93 | 0.0068 |
| Q32MZ4-3 | Isoform 3 of Leucine-rich repeat flightless-interacting protein 1 (LRRFIP1) | -1.98 | 0.0323 |
| P69905 | Hemoglobin subunit alpha (HBA1) | -1.99 | 0.0420 |
| P00918 | Carbonic anhydrase 2 (CA2) | -2.03 | 0.0420 |
| P02766 | Transthyretin (TTR) | -2.09 | 0.0049 |
| P01591 | Immunoglobulin J chain (IGJ) | -2.19 | 0.0137 |
| Q9BW30 | Tubulin polymerization-promoting protein family member 3 (TPPP3) | -2.35 | 0.0010 |
| P02765 | Alpha-2-HS-glycoprotein (AHSG) | -2.39 | 0.0186 |
| Q06830 | Peroxiredoxin-1 (PRDX1) | -2.43 | 0.0029 |
| P32119 | Peroxiredoxin-2 (PRDX2) | -2.48 | 0.0059 |
| Q16270 | Insulin-like growth factor-binding protein 7 (IGFBP7) | -2.52 | 0.0323 |
| Q13228-4 | Isoform 4 of Selenium-binding protein 1 (SELENBP1) | -2.54 | 0.0059 |
| P01620 | Ig kappa chain V-III region SIE (IGKV3-20) | -2.55 | 0.0440 |
| Q6P5S2 | UPF0762 protein C6orf58 (C6orf58) | -2.59 | 0.0059 |
| P15311 | Ezrin (EZR) | -2.64 | 0.0029 |
| O14745 | Na(+)/H(+) exchange regulatory cofactor NHE-RF1 (SLC9A3R1) | -2.67 | 0.0420 |
| P61769 | Beta-2-microglobulin (B2M) | -2.67 | 0.0137 |
| P68871 | Hemoglobin subunit beta (HBB) | -2.79 | 0.0420 |
| Q8N4F0 | BPI fold-containing family B member 2 (BPIFB2) | -2.80 | 0.0244 |
| P19823 | Inter-alpha-trypsin inhibitor heavy chain H2 (ITIH2) | -2.92 | 0.0420 |
| P23528 | Cofilin-1 (CFL1) | -2.96 | 0.0144 |
| P30838 | Aldehyde dehydrogenase, dimeric NADP-preferring (ALDH3A1) | -3.18 | 0.0059 |
| P07711 | Cathepsin L1 (CTSL) | -3.20 | 0.0323 |
| P10909-2 | Isoform 2 of Clusterin (CLU) | -3.27 | 0.0068 |
| Q14103 | Heterogeneous nuclear ribonucleoprotein D0 (HNRNPD) | -3.36 | 0.0244 |
| P13667 | Protein disulfide-isomerase A4 (PDIA4) | -4.04 | 0.0129 |
| P02042 | Hemoglobin subunit delta (HBD) | -4.32 | 0.0420 |
| O75347 | Tubulin-specific chaperone A (TBCA) | -4.63 | 0.0080 |
| P02545 | Prelamin-A/C (LMNA) | -4.92 | 0.0080 |
| P23141-2 | Isoform 2 of Liver carboxylesterase 1 (CES1) | -5.07 | 0.0059 |
| Q13938 | Calcyphosin (CAPS) | -6.11 | 0.0010 |
| P13010 | X-ray repair cross-complementing protein 5 (XRCC5) | -24.52 | 0.0248 |
| P08294 | Extracellular superoxide dismutase [Cu-Zn] (SOD3) | -24.87 | 0.0091 |
| Q5RHP9 | Glutamate-rich protein 3 (ERICH3) | -24.97 | 0.0092 |
| P27105 | Erythrocyte band 7 integral membrane protein (STOM) | -25.25 | 0.0323 |
| P09758 | Tumor-associated calcium signal transducer 2 (TACSTD2) | -25.30 | 0.0091 |
| P67936 | Tropomyosin alpha-4 chain (TPM4) | -25.46 | 0.0191 |
| Q15185 | Prostaglandin E synthase 3 (PTGES3) | -25.60 | 0.0244 |
| P34932 | Heat shock 70 kDa protein 4 (HSPA4) | -25.93 | 0.0020 |
| P50502 | Hsc70-interacting protein (ST13) | -25.99 | 0.0029 |
| Q96NY7 | Chloride intracellular channel protein 6 (CLIC6) | -26.08 | 0.0080 |
| P30043 | Flavin reductase (NADPH) (BLVRB) | -26.14 | 0.0092 |
| P06727 | Apolipoprotein A-IV (APOA4) | -26.16 | 0.0129 |
| P43353 | Aldehyde dehydrogenase family 3 member B1 (ALDH3B1) | -26.22 | 0.0010 |
| P30041 | Peroxiredoxin-6 (PRDX6) | -26.31 | 0.0092 |
| P07988 | Pulmonary surfactant-associated protein B (SFTPB) | -26.46 | 0.0330 |
| Q13740 | CD166 antigen (ALCAM) | -26.52 | 0.0059 |
| P19338 | Nucleolin (NCL) | -26.60 | 0.0244 |
| P05452 | Tetranectin (CLEC3B) | -27.06 | 0.0092 |
| P51884 | Lumican (LUM) | -27.13 | 0.0244 |
| P15328 | Folate receptor alpha (FOLR1) | -27.16 | 0.0440 |
| P13647 | Keratin, type II cytoskeletal 5 (KRT5) | -27.25 | 0.0080 |
| Q04917 | 14-3-3 protein eta (YWHAH) | -27.34 | 0.0323 |
| P02753 | Retinol-binding protein 4 (RBP4) | -27.42 | 0.0137 |
| P15090 | Fatty acid-binding protein, adipocyte (FABP4) | -27.70 | 0.0092 |
| P08727 | Keratin, type I cytoskeletal 19 (KRT19) | -27.95 | 0.0010 |
| P31946 | 14-3-3 protein beta/alpha (YWHAB) | -28.61 | 0.0029 |
| P00352 | Retinal dehydrogenase 1 (ALDH1A1) | -28.68 | 0.0059 |
| P06396 | Gelsolin (GSN) | -28.68 | 0.0080 |
| P05787-2 | Isoform 2 of Keratin, type II cytoskeletal 8 (KRT8) | -29.58 | 0.0092 |

<sup>a</sup>log<sub>2</sub>(FC) of VAP positive to Baseline, FC= fold change

**Table 3.** Comparison of ETA proteins between *Baselines* (from control and VAP patients) and *VAP positive*.

| Protein name | Gene ID | log2(FC)<sup>a</sup> | *p* value | Median Area Under Curve (AUC) |  |  |
| --- | --- | --- | --- | --- | --- | --- |
|  |  |  |  | Control<br>*Baseline* | VAP Patients |  |
|  |  |  |  |  | *Baseline* | *VAP positive* |
| Retinal dehydrogenase 1 | ALDH1A1 | -28.68 | 0.006 | 26.96 | 28.68 | 0.00 |
| Retinol-binding protein 4 | RBP4 | -27.42 | 0.014 | 28.44 | 27.42 | 0.00 |
| Tetranectin | CLEC3B | -27.06 | 0.009 | 25.23 | 27.06 | 0.00 |
| Isoform 2 of Keratin, type II cytoskeletal 8 | KRT8 | -29.58 | 0.009 | 27.59 | 29.58 | 0.00 |
| Gelsolin | GSN | -28.68 | 0.008 | 30.26 | 28.68 | 0.00 |
| Apolipoprotein A-IV | APOA4 | -26.16 | 0.013 | 25.07 | 26.16 | 0.00 |
| Pulmonary surfactant-associated protein B | SFTPB | -26.46 | 0.033 | 25.99 | 26.46 | 0.00 |
| Extracellular superoxide dismutase [Cu-Zn] | SOD3 | -24.88 | 0.009 | 24.53 | 24.88 | 0.00 |
| Keratin, type I cytoskeletal 19 | KRT19 | -27.95 | 0.001 | 27.81 | 27.95 | 0.00 |
| Tumor-associated calcium signal transducer 2 | TACSTD2 | -25.30 | 0.009 | 20.13 | 25.30 | 0.00 |
| X-ray repair cross-complementing protein 5 | XRCC5 | -24.52 | 0.025 | 21.24 | 24.52 | 0.00 |
| Keratin, type II cytoskeletal 5 | KRT5 | -27.25 | 0.008 | 25.77 | 27.25 | 0.00 |
| Fatty acid-binding protein, adipocyte | FABP4 | -27.70 | 0.009 | 28.34 | 27.70 | 0.00 |
| Folate receptor alpha | FOLR1 | -27.16 | 0.044 | 26.58 | 27.16 | 0.00 |
| Nucleolin | NCL | -26.60 | 0.024 | 24.80 | 26.60 | 0.00 |
| Peroxiredoxin-6 | PRDX6 | -26.31 | 0.009 | 25.34 | 26.31 | 0.00 |
| Flavin reductase (NADPH) | BLVRB | -26.14 | 0.009 | 25.21 | 26.14 | 0.00 |
| 14-3-3 protein beta/alpha | YWHAB | -28.61 | 0.003 | 27.98 | 28.61 | 0.00 |
| Heat shock 70 kDa protein 4 | HSPA4 | -25.93 | 0.002 | 25.93 | 25.93 | 0.00 |
| Hsc70-interacting protein | ST13 | -25.99 | 0.003 | 25.81 | 25.99 | 0.00 |
| Lumican | LUM | -27.13 | 0.024 | 25.53 | 27.13 | 0.00 |
| Tropomyosin alpha-4 chain | TPM4 | -25.46 | 0.019 | 26.43 | 25.46 | 0.00 |
| 14-3-3 protein eta | YWHAH | -27.34 | 0.032 | 26.10 | 27.34 | 0.00 |
| CD166 antigen | ALCAM | -26.52 | 0.006 | 26.37 | 26.52 | 0.00 |
| Glutamate-rich protein 3 | ERICH3 | -24.97 | 0.009 | 22.13 | 24.97 | 0.00 |
| Chloride intracellular channel protein 6 | CLIC6 | -26.08 | 0.008 | 25.22 | 26.08 | 0.00 |
| Peptidyl-prolyl cis-trans isomerase FKBP5 | FKBP5 | 1.71 | 0.032 | 0.00 | 23.39 | 25.10 |
| Ig kappa chain V-III region SIE | IGKV3-SIV | -2.56 | 0.044 | 0.00 | 28.12 | 25.57 |
| Ficolin-1 | FCN1 | 3.39 | 0.009 | 0.00 | 22.56 | 25.95 |
| Isoform 2 of Nucleoside diphosphate kinase A | NME1 | 26.56 | 0.029 | 0.00 | 0.00 | 26.56 |
| C-reactive protein | CRP | 28.11 | 0.037 | 0.00 | 0.00 | 28.11 |

<sup>a</sup>Log2(FC) of VAP positive vs. *Baseline* in VAP patients; *p*-value determined using Wilcoxon rank sum test.

**Table 4.** Reactome pathway analysis of differentially expressed proteins between Baseline and VAP positive.

| | Pathway | Significant proteins ($p < 0.05$) | $p$ value |
| --- | --- | --- | --- |
| Up Regulated | Neutrophil degranulation | FCN1;GGH;RETN;CFP;OLFM4;MPO;MMP9;NME1;CHIT1<br>PNP;S100A9;S100A8;VCL | 6.59E-14 |
|  | Innate immune system | FCN1;CRP;GGH;RETN;CFP;OLFM4;MPO;MMP9;NME1<br>CHIT1;PNP;MYH9;S100A9;S100A8;VCL | 6.24E-10 |
|  | Initial triggering of complement | FCN1;CRP;CFP | 9.27E-05 |
|  | Smooth muscle contraction | VCL;MYL12B | 6.68E-05 |
|  | EPH-Ephrin signaling | MYH9;MMP9;MYL12B | 5.55E-05 |
|  | RHO GTPases activate ROCKs, CIT, PAKS | MYH9;MYL12B | 9.89E-04 |
|  | Interconversion of nucleotide di- and triphosphates | AK2;NME1 | 6.07E-04 |
|  | Metal sequestration by antimicrobial proteins | S100A9;S100A8 | 3.20E-04 |
|  | Ficolins bind to repetitive carbohydrate structures on the target cell surface | FCN1 | 2.73E-04 |
|  | Metabolism of nucleotides | PNP;AK2;NME1 | 1.60E-03 |
| Down Regulated | Erythrocytes take up oxygen and release carbon dioxide | CA1;CA2; HBA1;HBA2;HBB | 4.33E-07 |
|  | Neutrophil degranulation | CD59;PRDX6;LGALS3;XRCC5;CFD;TTR;HSG;GSN;B2M;<br>ALDH2B1;HBB;STOM;CTSD;CTSH | 9.58E-06 |
|  | Chk1/Chk2(Cds1) mediated inactivation of Cyclin B:Cdk1 complex | YWHAB;YWHAH;YWHAZ | 1.02E-04 |
|  | Detoxification of reactive oxygen species | PRDX1;PRDX2;PRDX6;SOD3 | 2.06E-04 |
|  | TP53 regulates metabolic genes | PRDX1;PRDX2;YWHAB;YWHAZ;YWHAH | 4.92E-04 |
|  | Neurodegenerative diseases | PRDX1;PRDX2;LMNA | 6.77E-04 |
|  | Scavenging of heme from plasma | HBA1;HBA2;JCHAIN;KappaChainVII-SIE, KappChainC | 8.32E-04 |
|  | Amyloid fiber formation | TTR;GSN;B2M;APOA4 | 1.80E-03 |
|  | Platelet degranulation | LGALS3BP;CFL1;CFD;AHSG;CLEC3B | 2.55E-03 |

**Table 5.** Comparison of the ETA proteome among *Baseline*, *VAP positive* (day of diagnosis) and *pre-VAP* (2 days prior to diagnosis)

| Protein name | Gene ID | *VAP positive* |  | *pre-VAP* |  |
| --- | --- | --- | --- | --- | --- |
|  |  | log2(FC)<sup>a</sup> | *p* value | log2(FC)<sup>b</sup> | *p* value |
| Retinal dehydrogenase 1 | ALDH1A1 | -28.680 | 0.006 | -2.178 | 0.036 |
| Carbonic anhydrase 1 | CA1 | -1.854 | 0.024 | -1.375 | 0.022 |
| Hemoglobin subunit delta | HBD | -4.322 | 0.042 | -3.364 | 0.022 |
| Prelamin-A/C | LMNA | -4.922 | 0.008 | -1.703 | 0.036 |
| Alpha-2-HS-glycoprotein | AHSG | -2.389 | 0.019 | -1.022 | 0.022 |
| Isoform H7 of Myeloperoxidase | MPO | 1.206 | 0.019 | 1.759 | 0.022 |
| Keratin, type I cytoskeletal 19 | KRT19 | -27.954 | 0.001 | -2.935 | 0.022 |
| Pro-cathepsin H | CTSH | -1.644 | 0.019 | -1.539 | 0.035 |
| Tumor-associated calcium signal transducer 2 | TACSTD2 | -25.301 | 0.009 | -25.301 | 0.036 |
| Keratin, type II cytoskeletal 5 | KRT5 | -27.253 | 0.008 | -2.261 | 0.036 |
| Isoform 2 of Liver carboxylesterase 1 | CES1 | -5.067 | 0.006 | -1.794 | 0.036 |
| Peroxiredoxin-6 | PRDX6 | -26.307 | 0.009 | -26.307 | 0.036 |
| Peroxiredoxin-2 | PRDX2 | -2.481 | 0.006 | -2.402 | 0.022 |
| Aldehyde dehydrogenase family 3 member B1 | ALDH3B1 | -26.217 | 0.001 | -1.379 | 0.022 |
| Hsc70-interacting protein | ST13 | -25.990 | 0.003 | -25.990 | 0.022 |
| Adenylate kinase 2, mitochondrial | AK2 | 1.138 | 0.032 | 1.585 | 0.022 |
| Hemoglobin subunit beta | HBB | -2.790 | 0.042 | -2.657 | 0.022 |
| Hemoglobin subunit alpha | HBA1 | -1.992 | 0.042 | -1.792 | 0.022 |
| Isoform 4 of Selenium-binding protein 1 | SELENBP1 | -2.544 | 0.006 | -1.972 | 0.036 |
| Calcyphosin | CAPS | -6.112 | 0.001 | -1.790 | 0.022 |
| Isoform 3 of Leucine-rich repeat flightless-interacting protein 1 | LRRFIP1 | -1.980 | 0.032 | -1.150 | 0.036 |

<sup>a, b</sup> indicates log2(FC) of *VAP positive* to *Baseline* and *pre-VAP* to *Baseline*, respectively

To identify early VAP response mechanisms, we evaluated ETA collected two days prior to clinical diagnosis (*pre-VAP*). We identified 21 significantly differentially abundant proteins (*p <* 0.05) with a fold change >2 compared to *Baseline* (Figure 3C, Table 5). Consistent with our prior analysis, each of the 19 down-regulated proteins were associated with binding functions suggestive of scavenging and sequestration as an early host response to infection (Figure 4A-C). We observed recurrence of binding mechanisms and up-regulation of both isoform H7 of myeloperoxidase (MPO) and adenylate kinase 2 (AK2) in *pre-VAP* ETA (Figure 4C). While MPO has been involved in neutrophil mediated innate immunity (28), AK2 has been reported as a ubiquitous marker of cell lysis (29). This suggests their potential role in early defense against VAP infection, prompting us to explore neutrophil mediated pathogen processing. Longitudinal trends measured by LC-MS/MS of the two key components of neutrophil granules, MPO and ELANE, were confirmed by ELISA (Figure 4D). Both proteins were significantly higher in *pre-VAP* (4.8- to 5-fold compared to *Baseline*, adj. p < 0.044) and *VAP positive* (3.4 to 4-fold compared to *Baseline*, adj. p < 0.038), highlighting the rapid initiation of neutrophil degranulation and may serve as early detection markers.

**Figure 4.**
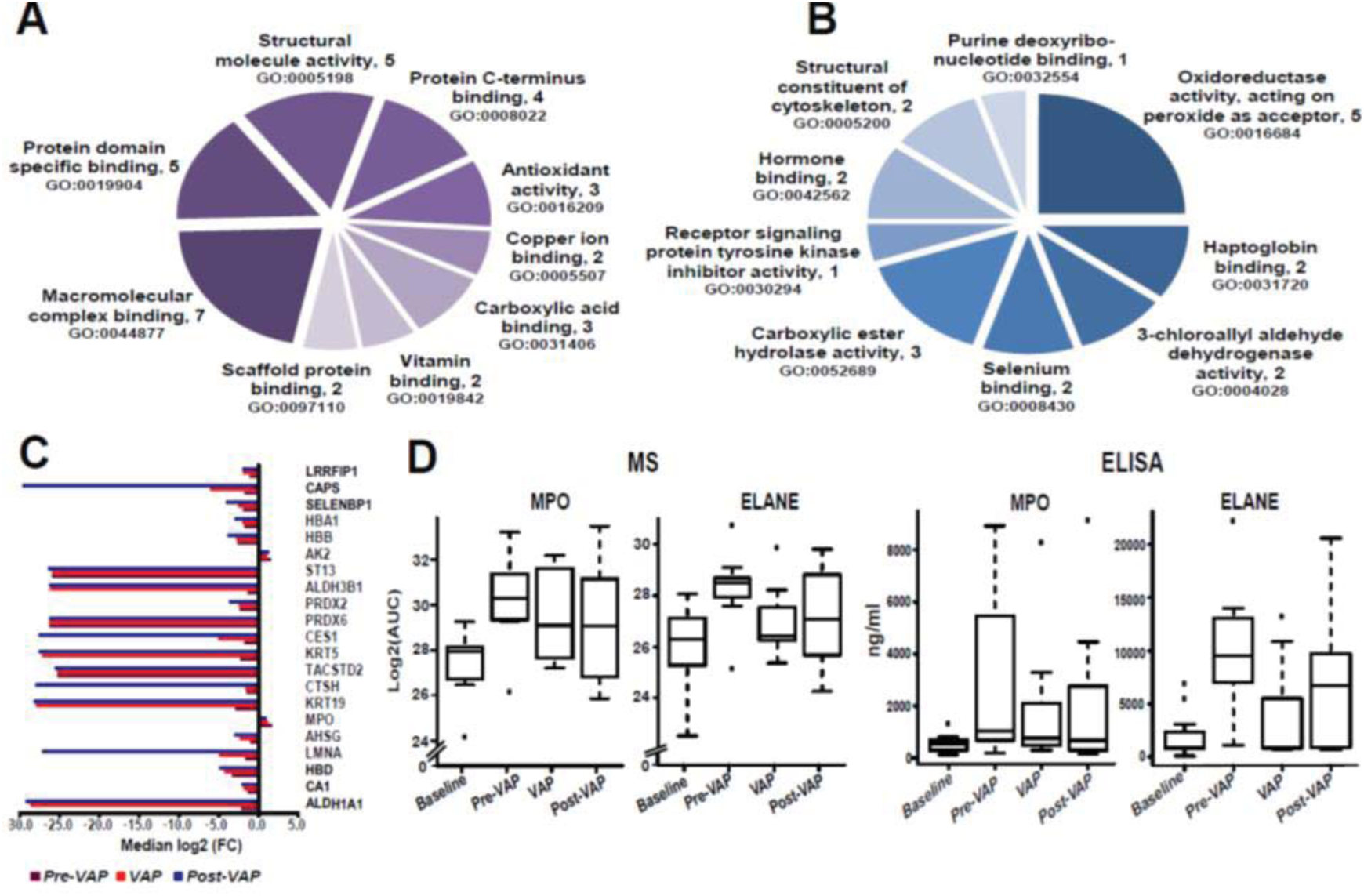
Analysis of functional enrichment and protein levels of differentially abundant proteins A. GO molecular function analysis of proteins only present in *Baseline* (at day of intubation). **B.** GO molecular function analysis of down-regulated proteins in *pre-VAP* and *VAP positive* relative to *Baseline*. **C.** Median log2(FC) of protein abundance for 21 significant proteins in *pre-VAP* (dark red), *VAP positive* (red), and *post-VAP* (blue) compared to *Baseline*. **D.** Measurement of MPO and ELANE levels in VAP patients using MS aided relative abundance, log2(AUC) (left) and ELISA (ng/mL of ETA) (right).

### VAP patients harbor metabolic signatures of oxidative stress

Similar to the proteomic analysis, Bland-Altman analysis showed no significant bias of metabolites in ETA and BAL matrices as the majority of data were within the limits of agreement for 95% confidence interval (mean difference of 0, upper limit of 4.7 and lower limit of −4.8) (Figure 5A). The unsupervised temporal clustering using median metabolite concentrations showed two distinct clusters of *Baseline* and post-*Baseline* samples. Unlike the temporal clustering of the ETA proteome, *Control* clustered more closely with *Post-VAP*. This suggests that at the time of intubation, ETA metabolomes of both control and VAP patients were similar and changed as infection progressed from *pre-VAP, VAP positive* to *post-VAP*. The clustering of *Control* with *post-VAP* may highlight an effect of residual inflammation due to intubation instead of infection (Figure 5B). We saw increased concentration of several metabolites at either *pre-VAP* or *VAP positive,* or in both. These metabolites were decreased in both *Control* and *Post-VAP*. These changes may reflect patient responses against infection. Further comparison of *VAP positive* and *Baseline* ETA identified a significant increase in 54 metabolites upon VAP infection (*p* < 0.05, fold change >2) (Figure 5B, Table 6). Of these, high concentrations of amino acids and C4-OH Pro in *VAP positive* ETA may have resulted from the activity of neutrophil proteases, matrix metalloproteinase-9 (MMP9) and ELANE, whereas increased concentration of Met-SO may indicate reactive oxygen species (ROS) formation by MPO, NADH oxidase or PNP during neutrophil degranulation (Figure 7C). In *VAP positive* ETA, we also observed a 2- to 5-fold increase of five polyamines as products of arginine catabolism, i.e. ADMA, ornithine, citrulline, spermine and spermidine. Elevated ADMA points towards ROS induced proteolysis of methylated proteins. Its inhibitory action on nitric oxide synthase (NOS) was measured as a decrease in Nitro-Tyr, also previously reported in community-acquired pneumonia (30). We detected increased acylcarnitines, glycerophospholipids and sphingolipids during VAP infection. Similar trends of plasma lipid metabolism have been reported in VAP patients (31).

**Figure 5.**
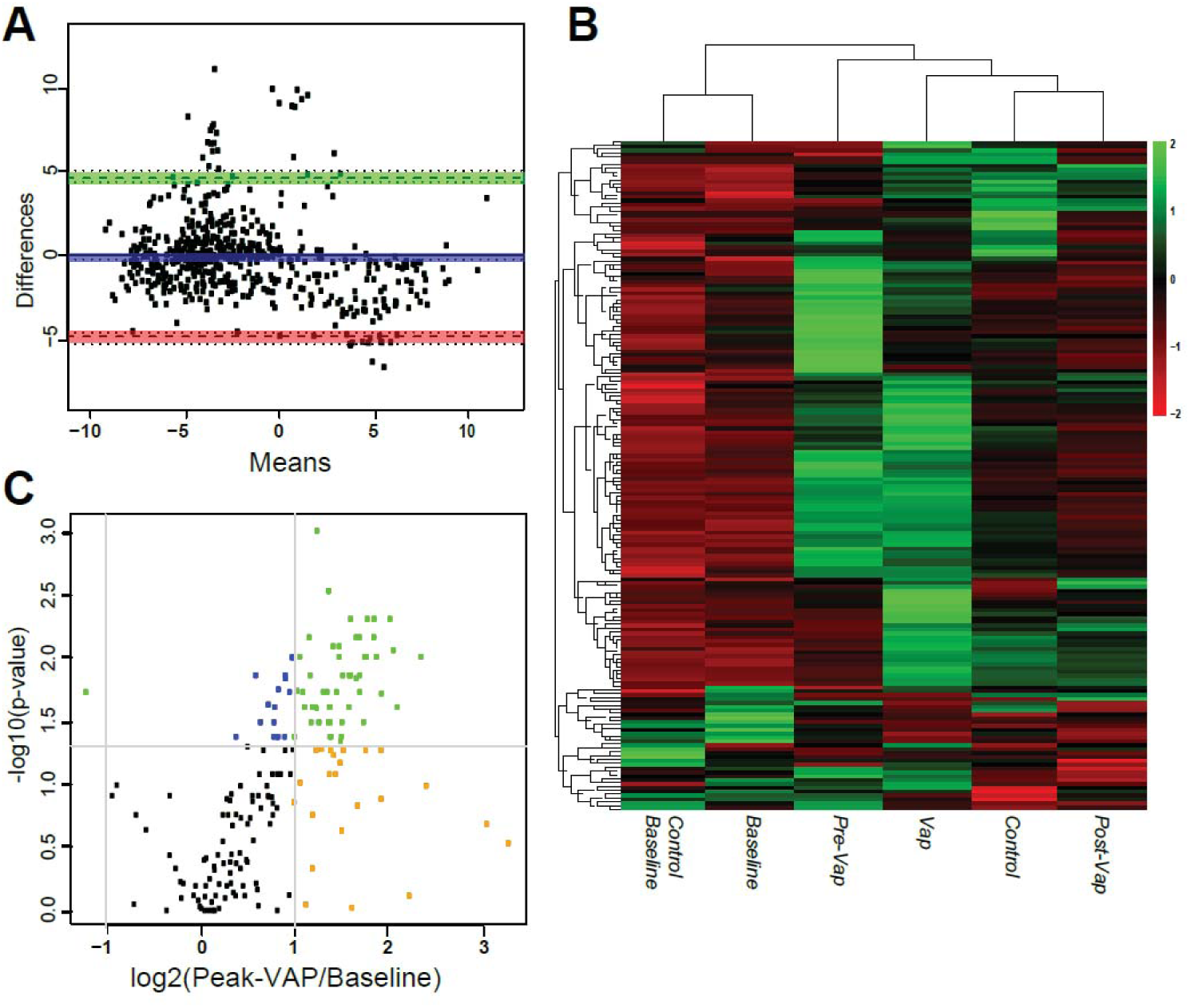
Differential metabolomic analysis A. Bland-Altman analysis of metabolite concentrations (μM) from ETA and BAL collected on the same day. **B.** Temporal clustering of clinical time points using median concentrations of metabolites in ETA. **C.** The volcano plot indicates differential concentrations of 185 metabolites in ETA by comparing the *VAP positive* to *Baseline*. (i) In blue, significant metabolites, (ii) in orange, metabolites for which the log2(FC) of metabolite concentration (*VAP positive* compared to *Baseline*) > 1 or < −1; (iii) in green, significant metabolites with log2(ratio) > 1 or < −1.

**Table 6.**
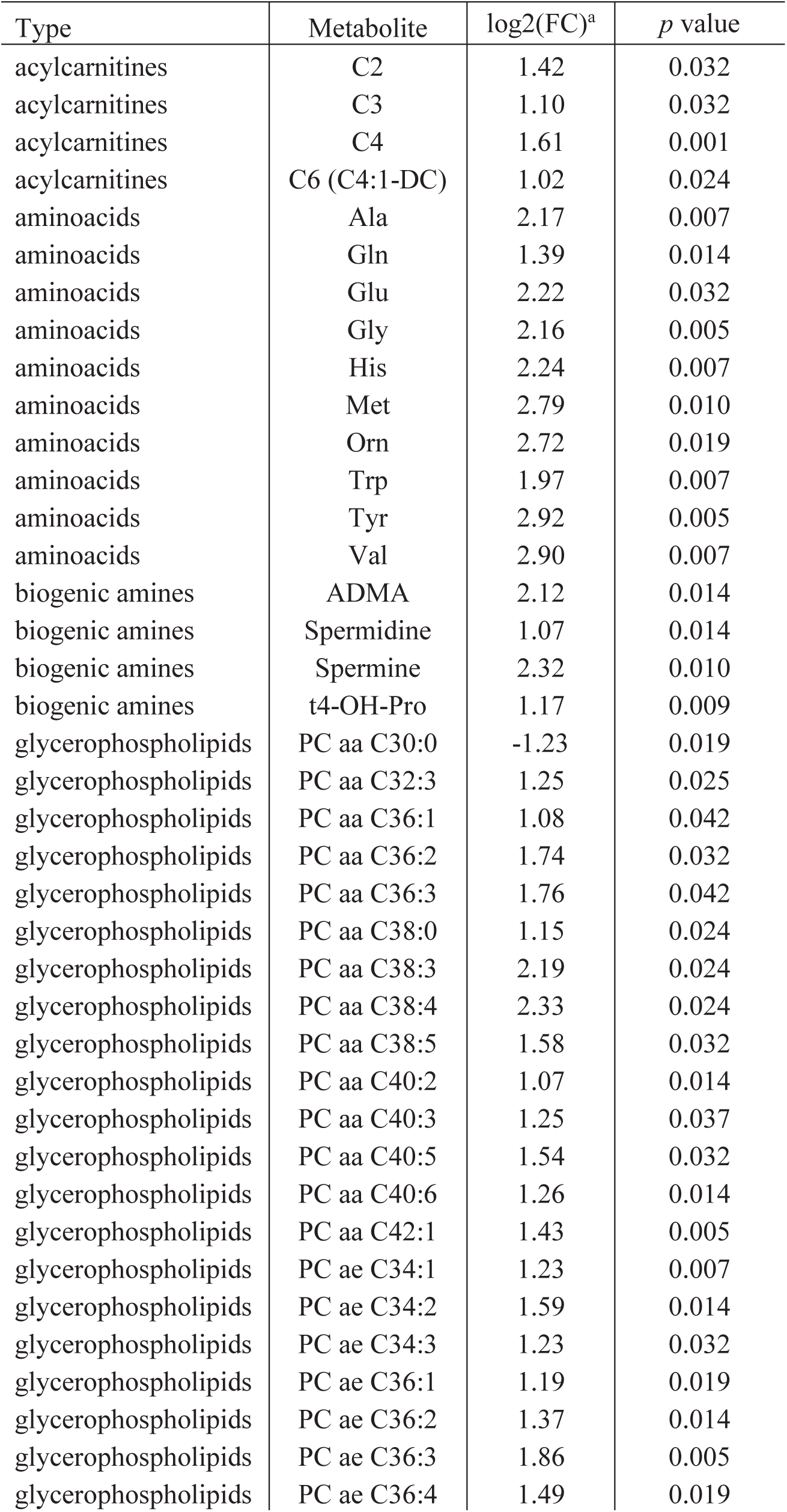
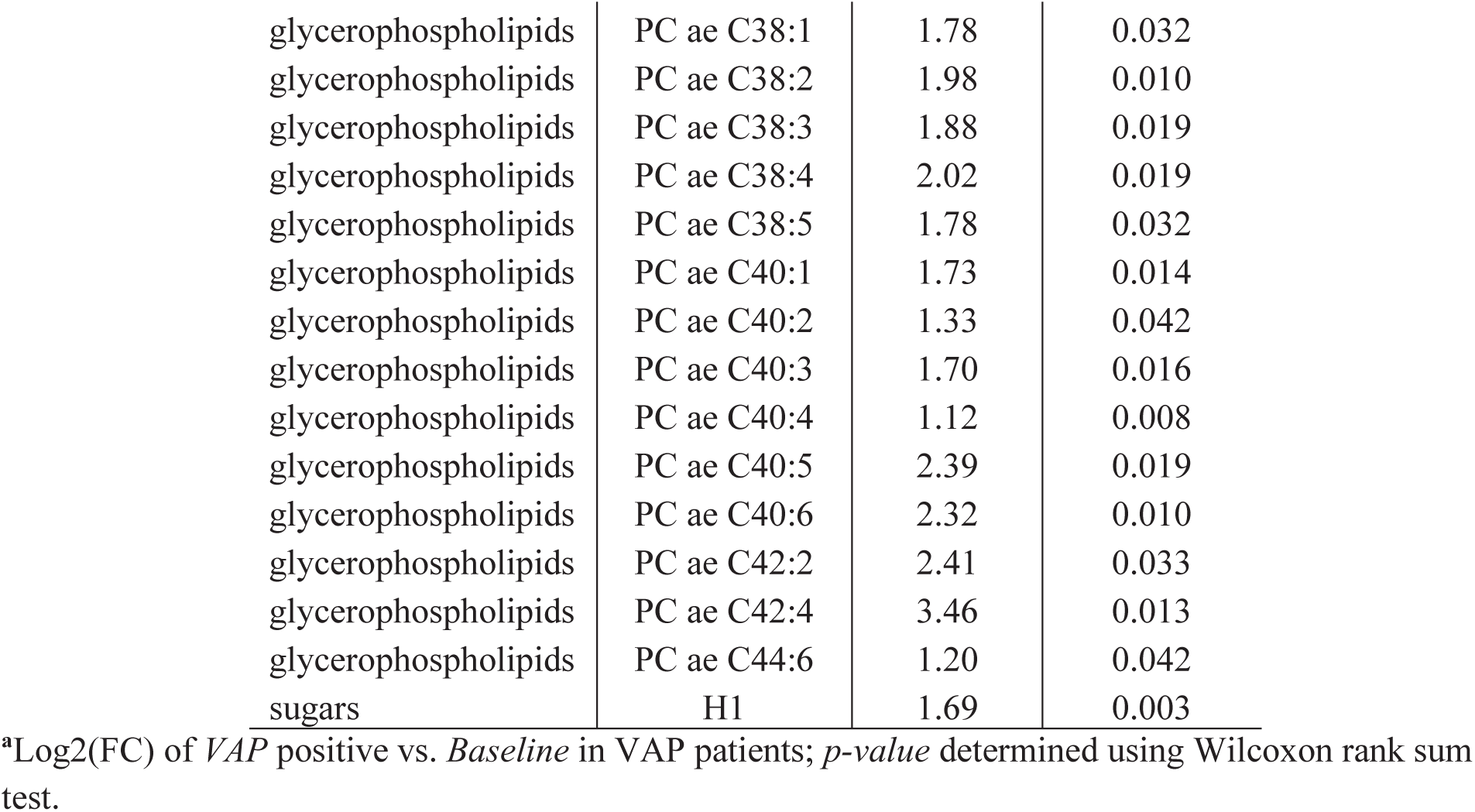
Differentially abundant metabolites between Baseline and the day of clinical diagnosis (VAP positive)

| Type | Metabolite | log<sub>2</sub>(FC)<sup>a</sup> | *p* value |
| --- | --- | --- | --- |
| acylcarnitines | C2 | 1.42 | 0.032 |
| acylcarnitines | C3 | 1.10 | 0.032 |
| acylcarnitines | C4 | 1.61 | 0.001 |
| acylcarnitines | C6 (C4:1-DC) | 1.02 | 0.024 |
| aminoacids | Ala | 2.17 | 0.007 |
| aminoacids | Gln | 1.39 | 0.014 |
| aminoacids | Glu | 2.22 | 0.032 |
| aminoacids | Gly | 2.16 | 0.005 |
| aminoacids | His | 2.24 | 0.007 |
| aminoacids | Met | 2.79 | 0.010 |
| aminoacids | Orn | 2.72 | 0.019 |
| aminoacids | Trp | 1.97 | 0.007 |
| aminoacids | Tyr | 2.92 | 0.005 |
| aminoacids | Val | 2.90 | 0.007 |
| biogenic amines | ADMA | 2.12 | 0.014 |
| biogenic amines | Spermidine | 1.07 | 0.014 |
| biogenic amines | Spermine | 2.32 | 0.010 |
| biogenic amines | t4-OH-Pro | 1.17 | 0.009 |
| glycerophospholipids | PC aa C30:0 | -1.23 | 0.019 |
| glycerophospholipids | PC aa C32:3 | 1.25 | 0.025 |
| glycerophospholipids | PC aa C36:1 | 1.08 | 0.042 |
| glycerophospholipids | PC aa C36:2 | 1.74 | 0.032 |
| glycerophospholipids | PC aa C36:3 | 1.76 | 0.042 |
| glycerophospholipids | PC aa C38:0 | 1.15 | 0.024 |
| glycerophospholipids | PC aa C38:3 | 2.19 | 0.024 |
| glycerophospholipids | PC aa C38:4 | 2.33 | 0.024 |
| glycerophospholipids | PC aa C38:5 | 1.58 | 0.032 |
| glycerophospholipids | PC aa C40:2 | 1.07 | 0.014 |
| glycerophospholipids | PC aa C40:3 | 1.25 | 0.037 |
| glycerophospholipids | PC aa C40:5 | 1.54 | 0.032 |
| glycerophospholipids | PC aa C40:6 | 1.26 | 0.014 |
| glycerophospholipids | PC aa C42:1 | 1.43 | 0.005 |
| glycerophospholipids | PC ae C34:1 | 1.23 | 0.007 |
| glycerophospholipids | PC ae C34:2 | 1.59 | 0.014 |
| glycerophospholipids | PC ae C34:3 | 1.23 | 0.032 |
| glycerophospholipids | PC ae C36:1 | 1.19 | 0.019 |
| glycerophospholipids | PC ae C36:2 | 1.37 | 0.014 |
| glycerophospholipids | PC ae C36:3 | 1.86 | 0.005 |
| glycerophospholipids | PC ae C36:4 | 1.49 | 0.019 |
| glycerophospholipids | PC ae C38:1 | 1.78 | 0.032 |
| glycerophospholipids | PC ae C38:2 | 1.98 | 0.010 |
| glycerophospholipids | PC ae C38:3 | 1.88 | 0.019 |
| glycerophospholipids | PC ae C38:4 | 2.02 | 0.019 |
| glycerophospholipids | PC ae C38:5 | 1.78 | 0.032 |
| glycerophospholipids | PC ae C40:1 | 1.73 | 0.014 |
| glycerophospholipids | PC ae C40:2 | 1.33 | 0.042 |
| glycerophospholipids | PC ae C40:3 | 1.70 | 0.016 |
| glycerophospholipids | PC ae C40:4 | 1.12 | 0.008 |
| glycerophospholipids | PC ae C40:5 | 2.39 | 0.019 |
| glycerophospholipids | PC ae C40:6 | 2.32 | 0.010 |
| glycerophospholipids | PC ae C42:2 | 2.41 | 0.033 |
| glycerophospholipids | PC ae C42:4 | 3.46 | 0.013 |
| glycerophospholipids | PC ae C44:6 | 1.20 | 0.042 |
| sugars | H1 | 1.69 | 0.003 |

<sup>a</sup>Log<sub>2</sub>(FC) of VAP positive vs. *Baseline* in VAP patients; *p-value* determined using Wilcoxon rank sum test.

### VAP pathogen specific peptide signatures are found in ETA

Using meta-proteomics, we attempted to identify species specific signatures of VAP pathogen (Supplementary Table S4) in our study samples. We identified a total of 137 peptides corresponding to 124 proteins specific to VAP pathogens derived from the literature (Figure 6A, Table 7, Supplemental Table 4). Of these, 40 peptides were linked to 4 gram-positive VAP species, 95 peptides to 10 gram-negative VAP species, and 2 peptides were linked to *Candida albicans*. Out of 40 Gram-positive specific peptides, 35 were specific to *Propinibacterium*, *Staphylococcus*, *Streptococcus* or *Enterococcus* genera. Furthermore, 59 peptides were specific to Gram-negative genera and 21 were explicit to *Escherichia coli, Enterobacter clocae, Klebsiella aerogenes, Pseudomonas aeruginosa* and *Serratia marcescens*. Our metaproteomic approach using species-specific peptide signatures confirmed the culture-based diagnosis in 13 out of 16 VAP patients (approximately 80%). In control patients, low peptide counts specific to gram-negative pathogens were detected in a majority of time points, whereas peptides attributed to gram-positive were only detected at day 1 post intubation in two patients (Figure 6B). Based on peptide specificity, gram-negative pathogens were detected in 9 out of 10 VAP patients at all-time points. Gram-positives were only detected in 4 out of 10 patients, with highest peptide counts in patient 16. This suggests that VAP infection is majorly dominated by gram-negative bacterial species. Furthermore, species-specific peptides were detectable days before culture based diagnosis.

**Figure 6.**
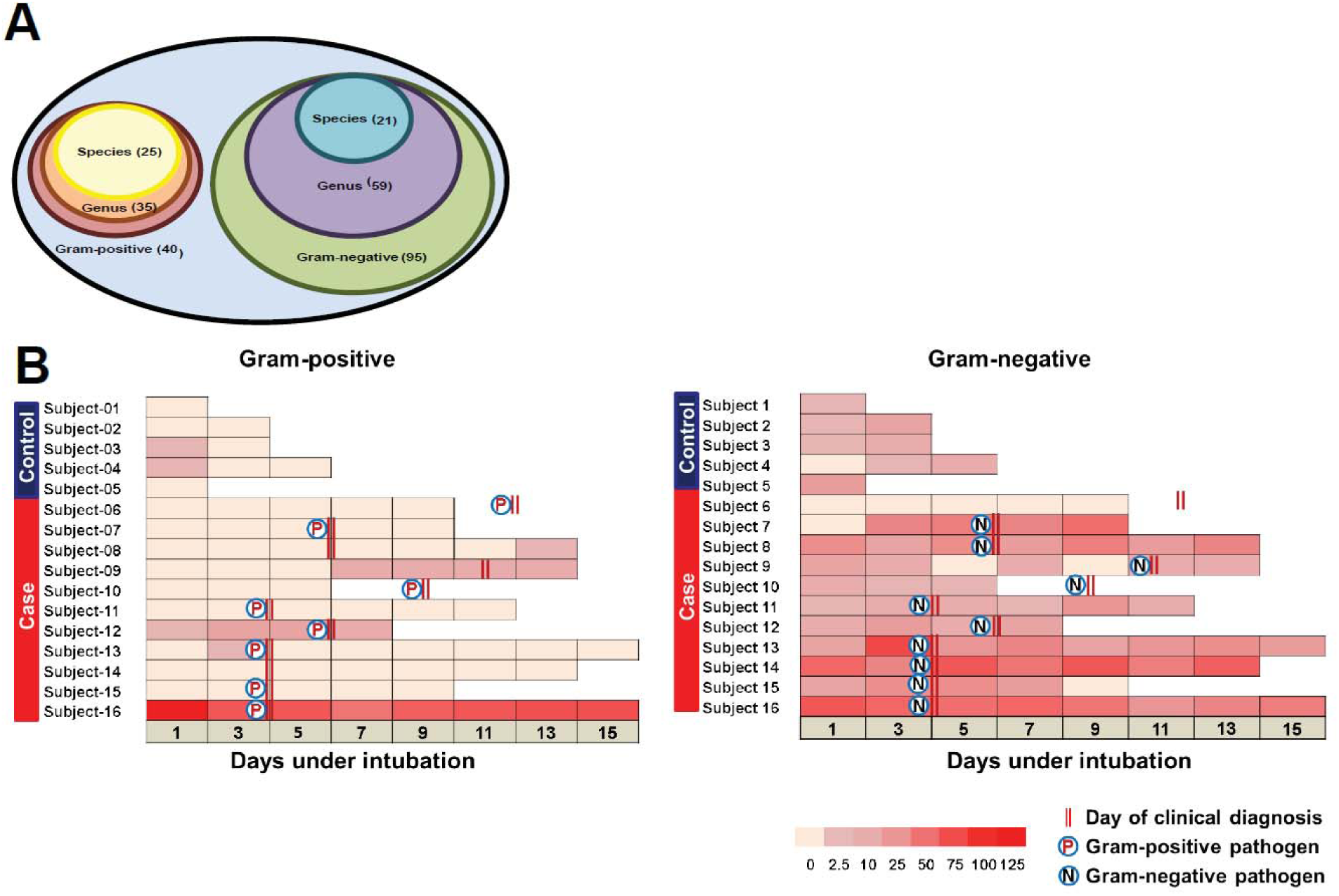
Pathogen peptide signatures in ETA A. VAP pathogen specific peptides detected in ETA. In gram-positive group (40), genus specific (35) and species (25); gram-negative group (95), genus specific (59) and species (21) specific peptides were identified. **B.** Distribution of peptides specific to gram-positive and gram-negative bacterial pathogens in control and VAP patients. A color gradient from light to bright red corresponds to number of peptides detected during each intubation time point. P and N indicate gram-positive and gram-negative pathogens detected using culture test, respectively. Day of clinical diagnosis is shown as parallel red lines.

**Table 7.** Species-specific peptides leading to identification of bacterial proteins in ETA.

| Peptide Sequence | Protein | Specificity | Gram nature |
| --- | --- | --- | --- |
| MQGAVWGIDSFDQWGVELGK<br>FDPTFSALKSR<br>TGLNEGILISDGPLPTR | JCM18916_1830<br>JCM18909_2989<br>HMPREF0675_5076 | *Propinibacterium acnes* | Pos |
| NALIIEDTGDNNVVK<br>SVVNVQPTNEENKK<br>ELIEASMTYSDNTANNK<br>GTFIIDPDGVVQASEINADGIGR<br>IIEQIGTYNPTSANAPEIK<br>IVDYPGKEDIAVVQVEEK<br>LANGGENSIEGTANLNELSNNSNNKPINSDSVK<br>MIAPVLEELAADYEGK<br>NLFTGWEDVNLSEQGINEATR<br>NNIADKNVETLNEK<br>VDTAQVNGDTR<br>EANIIDKTDNENVK<br>KDGMLLFNLHQGALLQGMTATQLLLL VFFSSTFTACSVTMTMLLK<br>KYKDMTQEEVK<br>LAYEINDFKDGFYNIVR<br>NGDSVAQSYDMIDNAPEEK<br>VGEGVTSGGENTLTK<br>YLGDEEISVSELK<br>ATLPNEDVVESDPSTTK | greA<br>sdrD<br>blaZ<br>ahpC<br>rpsP<br>SAKOR_01758<br>ear<br>SAKOR_01066<br>pgm_2<br>SAUSA300_pUSA010004<br>HMPREF0776_2043<br>SAMEA3448920_01277<br>ST398NM01_2633<br>A7U48_1386<br>rpsF<br>tufA<br>SAMSHR1132_16580<br>fusA<br>ST398NM01_1474 | *Staphylococcus aureus* |  |
| EMIVPDQIIMGLILYAGDAK<br>MIQPFIFLENNK | chbA<br>ERS020507_01391 | *Streptococcus pneumoniae* |  |
| MGANDNILKGYSTFMMEMLQCKNIISMMTTR | Cd36_22850 | *Candida dubliniensis* |  |
| LAGGFAYVEVLPK<br>QFPAMTQTMIEGLR<br>SERELEMSTEIAR<br>TLTRESLAIP<br>ADAVLHDTNPLYFIK | rfaC<br>nohA2<br>A1WK<br>ACU57_03400<br>glnH | *E.coli* | Neg |
| AFWDVVNWDEAAAR<br>GLKDDIKALQK<br>SGSSYNPLYNLMFQAGVPPRYMR<br>VCAGIVGAALLLLAGCSSNELSAAGQGVR<br>WATDLNSVIDNVLR | sodA<br>pilX5<br>ASV07<br>A3461<br>yihY | *Enterobacter cloacae* |  |
| QTRPGKNEQLFFMYK | EHR_08925 | *Enterococcus hirae* |  |
| EFLENYLMTDPGLEAVNNDKPLGAVALK<br>LYTTNADGELLTIDTDSNK<br>QNQLQLDEAASQLLCYCYEGNLLALAQALERLSLLWPDGK<br>VDMDASAALK<br>YLFECQQGDGNAPK | malE<br>EAE<br>holA<br>ASV18<br>zinT | *Klebsiella aerogenes* |  |
| EMMSILELANTDAGSGSGGEELASER<br>HWHGAAPSTAMMHIAVTATLDGKNVNWMEK<br>MNALRGWLAALGSMLLASAAQLGMR<br>NILSNINWGWR | PAERUG_E15_London_28_<br>01_14_03978<br>PAMH27<br>arnF<br>PAMH27_3286 | *Pseudomonas aeruginosa* |  |
| MSLTINGITVPDSAMAR<br>YLFECQDAGSLAPK | BHU62_19855<br>zinT | *Serratia marcescens* |  |

Pos=Gram-positive, Neg=Gram-negative

## Discussion

The present study describes VAP mediated host responses in ETA and BAL of 16 intubated patients. This is also the first detailed characterization of the ETA proteome and metabolome. We detected 3067 unique proteins in ETA sampled longitudinally, compared to 1139 proteins in BAL. We also observed >10% increase in unique BAL proteins compared to previous studies (16, 32). Despite our observation of a 3-fold higher proteome diversity and although less invasive, ETA has been historically overlooked in favor of BAL (33). Our study revealed that ETA is functionally diverse and highly enriched in proteins involved in innate and adaptive immunity, suggesting that it is an attractive source to study lung infection. In VAP patients, we observed up-regulation of inflammation and neutrophil-mediated innate immunity in the respiratory tract during VAP infection, leading to pathogen processing (Figure 7). Elevated levels of vinculin and myosins may imply extracellular matrix (ECM) adhesion and migration of neutrophils (Figure 7B) (34, 35). Increased abundance of pathogen recognition molecules ficolin-1 and properdin in ETA may be linked to complement system activation via interaction with bacterial polysaccharides (36). In support of this, antimicrobial neutrophil proteins S100A8 and S100A9, with proinflammatory and chemotactic activity (37), were highly elevated during VAP infection. Furthermore, the VAP-associated increase in granule proteins, such as chitotriosidase-1, gamma-glutamyl hydrolase, resistin, olfactomedin-4, and MMP9 (38, 39) all point towards inflammation and neutrophil degranulation to promote pathogen clearance (Figure 7B). Li and colleagues predicated an increase in plasma MMP9 with the severity and progression of VAP infection (40). During infection, release of these hydrolytic granule proteins upon neutrophil degranulation may have detrimental effect on ECM of airway epithelium. The high concentrations of free amino acids as well C4-OH Pro in ETA suggest altered ECM integrity through collagen degradation. Effect of neutrophil metalloproteases on ECM modulation has been previously studied (41). The neutrophils are key players of pathological inflammation in lung infections such as VAP (42). In our study, we observed increased lung inflammation and neutrophil degranulation during VAP infections which may promote formation and release of neutrophil extracellular traps (NETs). The role of NETs in VAP pathogenesis has been recently investigated in BAL of 100 critically ill patients (43).

**Figure 7.**
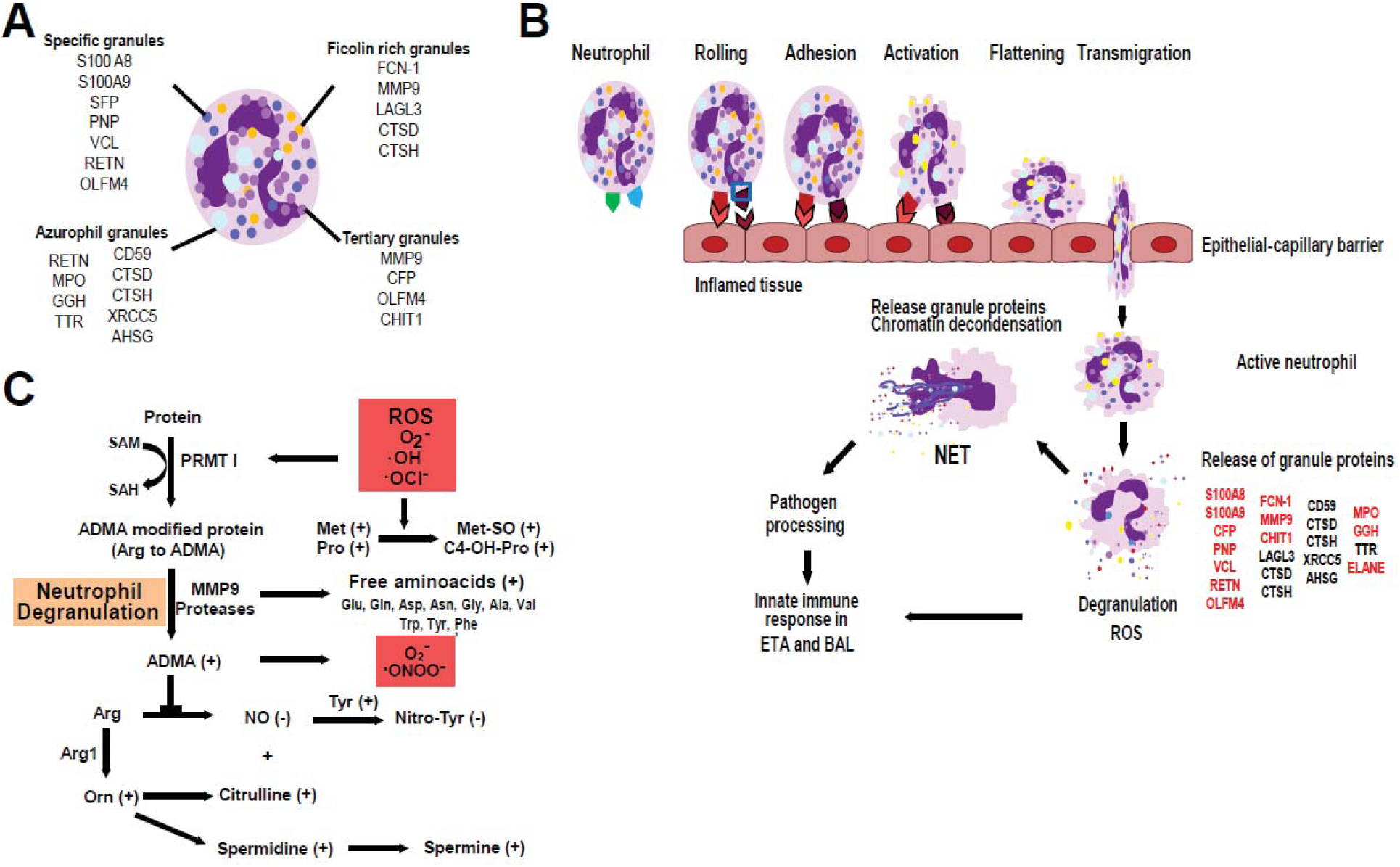
Proposed mechanisms of neutrophil-mediated innate immune responses in the respiratory tract upon VAP infection A. Neutrophil granules and their proteins detected in ETA and BAL. **B.** Recruitment of neutrophils at the site of inflamed trachea and alveoli in response to pathogen stimuli. Neutrophil rolling, adhesion and transmigration into inflamed alveoli and tracheal tissues are facilitated by VCL, ACTN1, MYYH9, and MYH12. Activation of neutrophils can induce degranulation and release granule proteases, oxidases, peroxidases and other antimicrobial proteins for pathogen processing (51). **C.** Proposed metabolic fate of neutrophil degranulation and reactive oxygen species (ROS)-induced oxidative stress in the respiratory tract during VAP infection.

Although, NETosis provides a defense network against pathogen infection, studies have shown that exaggerated NETs can be detrimental to the lung environment (44). Considering the tissue protective effect of polyamines (45), their high levels at VAP infection may have role in tissue protection as a balancing mechanism against adverse impact of neutrophil degranulation on lung epithelium.

Overall, ETA investigation suggests a cascade of neutrophil degranulation events starting with neutrophil recruitment, adhesion and migration towards the site of infection. Induction of oxidative stress activates degranulation and chromatin decondensation as a host response against VAP. Validation of elevated ELANE and MPO by ELISA confirmed the role of neutrophil degranulation in early host response to VAP. Studies have demonstrated higher specificity (93 to 96%) and sensitivity (75 to 91%) of serum CRP at levels ranging from 48 to 200 mg/L in diagnosing pneumonia (46, 47). The presence and role of CRP and NME1-2 only at *VAP positive* is still unclear and warrants further investigation.

We also observed the modulation of protein catabolism, ROS synthesis, polyamine and lipid metabolism in VAP ETA. The formation of ROS was measured using protein inducers (MPO and purine nucleoside phosphorylase) and was confirmed by metabolic indicators (Met-SO, C4-OH-Pro and ADMA) (28, 48, 49). Metzler et al. have further shown ROS-triggered translocation of ELANE to the nucleus and subsequent induction of chromatin decondensation. Oxidative stress is characteristic of neutrophil degranulation and has been reported in other pulmonary diseases (50). Our hypothesis is corroborated through the investigation of longitudinal ETA sampling, which has identified host innate immune mechanism of pathogen processing and provided enhanced granularity of VAP progression Through meta-proteomics analysis, we demonstrated that ETA and BAL harbored pathogen protein products specific to VAP pathogens. Since microbial peptide abundance was low, quantitation was beyond the scope of this study. The utility of these species specific peptides in VAP diagnosis will be explored in future.

Intubation is one of the most common interventions in critical care and has been linked to increased susceptibility of lung infection and mortality. Intubation procedure, length of stay and inappropriate antibiotic treatment, as well as pre-existing conditions such as compromised or weakened immunity may contribute to microbial dysbiosis and development of pneumonia. Ultimately, these alterations to the host environment may be reflected at the pulmonary interface. Our study has focused on ETA to conduct a molecular survey of upper airways during VAP development and progression. We have shown that ETA is reflective of a rich and diverse airway proteome. In VAP, we identified an early upregulation of immune-modulatory proteins associated with an early host response to infection. We also looked for VAP pathogen peptides in ETA, and detected unique, species-specific peptides correlated with cultures. In the majority of VAP patients, these distinctive pathogen signatures were present at least 1 to 2 days earlier than the BAL culture based diagnosis. Although presenting distinctive features from BAL, ETA may be an attractive alternative for earlier and cost effective clinical diagnosis of pneumonia in intubated patients.

## Acknowledgements

We thank the patients, their relatives and the medical staff at HonorHealth for their contributions and support towards this project. We thank Nancy Linford, PhD (Linford Biomedical Communications, LLC, Washington, USA), for reviewing this manuscript. We thank the Flinn Foundation for funding the sample collection (Grant ID#1975). This study was funded by institutional funds at TGen.

## Data Availability

The mass spectrometry proteomics data have been submitted to ProteomeXchange Consortium via the PRIDE partner repository with dataset identifier PXD010715. Metabolites quantified in this assay are available in Supplemental Table S1.

## Author contributions

PP, EM and FZ designed the study; CKH and KL performed sample collection and provided cohort clinical details; KVP, MIM have equally contributed in designing and performing experiments; KGM performed statistical analysis; KVP, MIM, KGM and PP interpreted results; YY and ZE guided the statistical analysis and reviewed the data interpretation, KVP, MIM, KGM and PP wrote the manuscript; all authors read and approved the final manuscript.

